# DataPLAN: a web-based data management plan generator for the plant sciences

**DOI:** 10.1101/2023.07.07.548147

**Authors:** Xiao-Ran Zhou, Sebastian Beier, Dominik Brilhaus, Cristina Martins Rodrigues, Timo Mühlhaus, Dirk von Suchodoletz, Richard M. Twyman, Björn Usadel, Angela Kranz

**Author notes:** Correspondence: X.-R. Z.; /61-85881, A.K.; /61-85833.

## Abstract

Research Data Management (RDM) is a system for the effective handling of research data that enables scientists to structure their research questions and ensure best practices throughout the data lifecycle, from acquisition, computation and annotation to data publication and re-use. Data management plans (DMPs) are documents that formally set out the RDM of a project and are required by many funding bodies. DMPs help to organize and structure RDM strategies, thus promoting data findability, accessibility, interoperability and reusability (FAIR). Although DMPs incorporate methods and standards that can be reused by different research projects, the standardization of DMP content is not as evident as the standardization of RDM practices and data/metadata. To address this issue in the plant sciences, we developed DataPLAN – a tool that combines a questionnaire with pre-written standardized responses. We wrapped the questionnaire in a serverless single-page web application that can then generate standardized responses from DMP templates. The current templates cater to plant research grant proposals for Horizon 2020, Horizon Europe and the German Research Foundation (Deutsche Forschungsgemeinschaft, DFG). In the future the range of templates will be extended to accommodate other funding schemes, thereby enabling more users to generate their own templates. The DataPLAN web application is open-source and does not require an internet connection. By utilizing DataPLAN, the workload associated with creating, updating, and adhering to DMPs is significantly reduced.

## Introduction

A data management plan (DMP) documents the methods, procedures, and policies for handling, sharing and analyzing research data throughout a project’s life cycle and beyond [1,2]. Its purpose is to set out a clear and structured approach to manage all types of anticipated research data effectively, including strategies for data acquisition, organization, documentation, storage, access, and sharing [3,4]. The DMP therefore contributes to efficient and productive research data management (RDM) and to the FAIRness of the data, making it findable, accessible, interoperable, and reusable [4]. Many funding bodies require a DMP, and in such cases a draft DMP should be defined in the proposal, then formalized during the project (before data/metadata collection begins), thus encompassing the entire RDM strategy of the project (**Figure 1a**).

**Figure 1.**
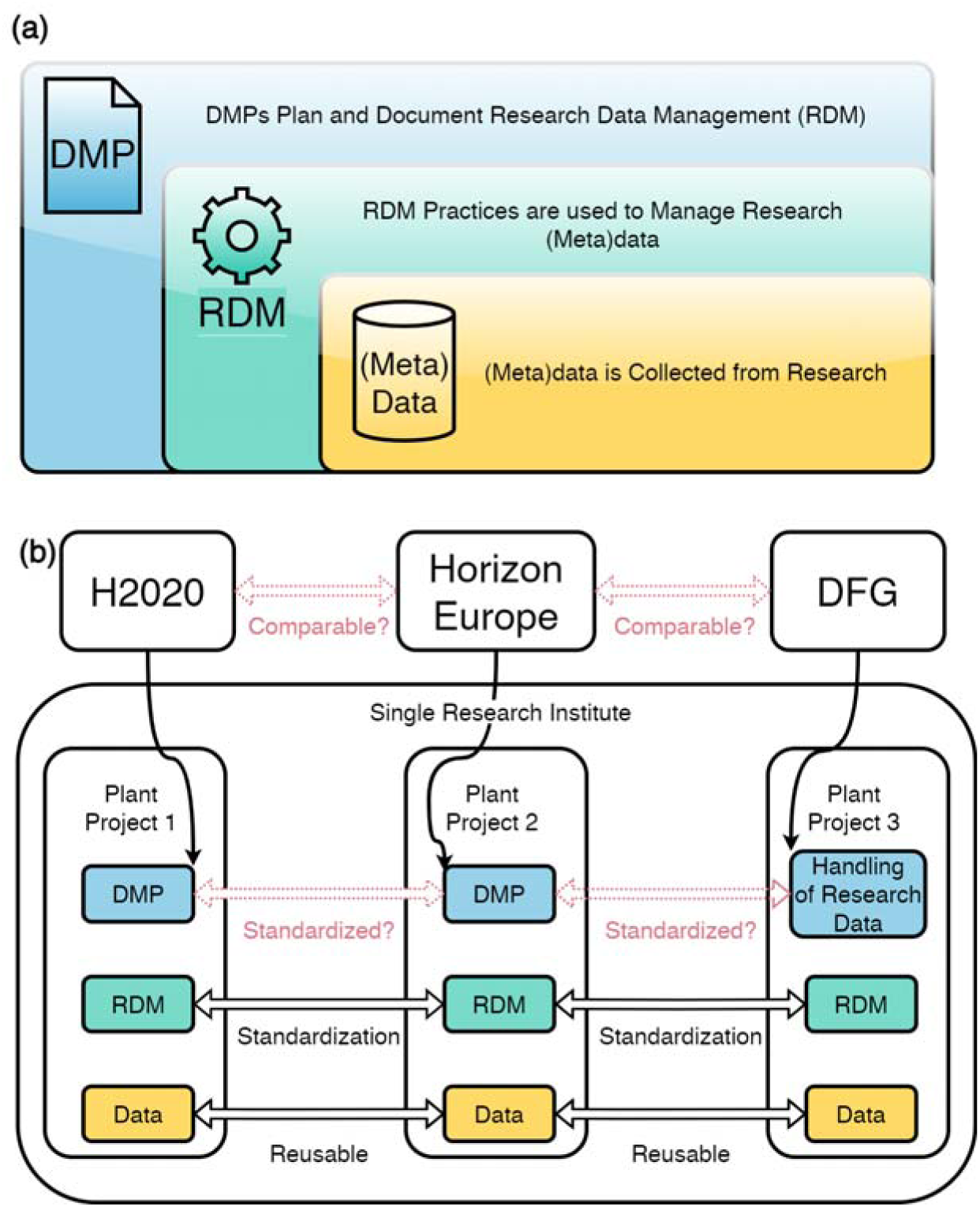
A standardized DMP is enabled by standardized RDM and reusable metadata/data. (a) DMPs document RDM practices for data/metadata; (b) Although DMPs encompass reusable standardized RDM practices [1], the standardization of the DMP content is not as evident as the standardization of RDM practices and data/metadata between projects [5,6].

The need for effective RDM has increased by factors such as the growing volume and complexity of research data, the requirements set by funding bodies such as Horizon 2020, Horizon Europe and the German Research Foundation (Deutsche Forschungsgemeinschaft, DFG) [7–9], and the need for data reuse and reproducibility. Prominent funding bodies, such as Horizon Europe [10], the DFG [11], the National Institute of Health (NIH) [12], the National Science Foundation (NSF) in the United States [13] and Swiss National Science Foundation (SNSF) [14] expect applicants to submit RDM related documents together with the proposal. The checklist of DFG’s “Handling of Research Data”, which is treated as a DMP equivalent by multiple universities [15,16], consists of 19 questions that cover various aspects of data management, whereas the Horizon Europe DMP questionnaire is more extensive, comprising 45 questions. By answering these questions, applicants are expected to demonstrate their understanding of good RDM practices and their commitment to implementing them. Although DMPs serve an essential purpose to document RDM practices within plant science projects, the standardization of DMP content is not always as evident as the standardization of RDM and data/metadata between projects. For example, consider three projects proposed by the same partners (Projects 1-3 in **Figure 1b**) that share infrastructure and protocols, but focus on different plant species within the same family. Ideally, their RDM measures should be reusable and consistent. However, if Project 1 receives funding from Horizon 2020, Project 2 is submitted to Horizon Europe, and Project 3 to the DFG, their DMPs and “Handling of Research Data” will vary in structure and content. This fragmented approach to DMPs leads to a lack of consistency and uniformity, posing challenges to the FAIRness of the DMPs and the data/metadata they manage. First, the findability of information becomes more difficult because the connection between the three projects becomes less apparent, the lack of a unified structure and standardized approach across DMPs can hinder the identification and retrieval of relevant information. Second, maintaining interoperability becomes challenging due to the differing structures and formats of the DMPs. Text comparison tools such as *git diff* and other methods that reveal changes in code become less effective when applied to DMPs with distinct structures. This makes it harder to track and understand the modifications made to the DMPs over time. Finally, the inability to replicate RDM content across different DMPs results in limited reusability. Despite similarities in RDM measures among the projects, the fragmented nature of the DMPs makes it difficult to transfer and adapt content efficiently from one DMP to another. Overall, the lack of consistency and uniformity across funding bodies can limit the FAIRness of the DMPs, leading to inefficient RDM practices. Efforts to promote greater harmonization and standardization of DMP requirements and structures could address these challenges and enhance the overall FAIRness and utility of DMPs across diverse projects.

Standardized DMPs need to incorporate decentralized RDM practices [17], because each research field may have its own set of standards, protocols, and best practices for data management [18–24], and RDM practices are often tailored to the specific requirements and characteristics of different disciplines. An overgeneralized approach to data management is ineffective, because different research fields face unique data management challenges and opportunities [6]. Therefore, good DMPs must achieve a balance between standardization and customization. A DMP for an international project must also consider the increasingly onerous legal obligations for data management. The number of laws and directives that govern how research data should be managed, shared, and accessed is increasing, especially in the context of international collaborative projects that involve cross-border data transfer. For example, under EU legislation, the utilization of genetic resources requires permission from the country of origin [25,26]. These requirements and standards can vary depending on the country, region, and research topic.

Multiple tools have been developed to tackle the challenges described above [2,27–38]. These tools help researchers to write a comprehensive DMP by providing a structured framework and guidelines. However, they vary in their support for the preparation of DMPs for specific funding bodies [2,27,28,39], the ability to focus on domain-specific RDM approaches [35,37,40], and the inclusion of relevant laws, directives and regulations [2,27,28,33,35,36,41,42]. To our knowledge, there is no DMP tool available that fully satisfies the needs of the plant research community.

The volume of research data in the plant sciences is growing exponentially [43,44]. Technologies such as nanopore sequencing [45–47] and high-throughput phenotyping [48,49] have increased the rate of data collection. The analysis of such data will provide a deeper understanding of plants leading to the generation of new knowledge and new paths to exploitation. To handle this data explosion, DMPs can be used to plan new research endeavors but they are tedious to prepare from the blank page. Many research methods associated with the fundamental plant sciences, such as genome sequencing and assembly or phenotyping experiments follow a standardized workflow and vary only to a certain amount with respect to the plant species. The standardized plant research pipelines would enable the standardization of DMPs, which would significantly speed up the funding application process and improve the quality of RDM related to plant fundamental sciences.

The quality of RDM and in a narrower sense the content and focus of a DMP can be improved by using data management platforms and infrastructures. The fundamental plant sciences deal with the characterization of plants, for example plant growth, crop yields and biomass production. This involves diverse data types including omics data and imaging datasets, which must be managed, integrated and annotated effectively to ensure the meaningful interpretation of results. As part of the German National Research Data Initiative (NFDI), the DataPLANT consortium [50] provides a research data infrastructure for the plant sciences with a user centric approach to RDM practices.

Here we present DataPLAN, a web-based and easy-to-use DMP generator tool that is part of the DataPLANT services framework. DataPLAN creates standardized templates for the generation of DMPs that currently fulfills the requirements of three funding bodies: DFG [11], Horizon 2020 [51] and Horizon Europe [10]. DataPLAN also provides a practical user guide to facilitate RDM approaches. DataPLAN’s source code is freely available on GitHub under a GPL-3.0 license (https://github.com/nfdi4plants/dataplan; accessed on 25 June 2023).

## Results

### User Interface

#### Main menu

The DataPLAN user interface (UI) features a straightforward design with a live preview on the left and a questionnaire on the right of the window (**Figure 2**). Users can navigate and interact with the tool seamlessly using the same webpage throughout the session. The interactive UI provides flexibility and offers users content without leaving the main interface. The main menu above the live preview has four dropdown menus: *Templates, Import, Export*, and *Help*. These provide 22 clickable options. Clicking the *Templates* button enables users to switch between the three current templates (H2020 DMP, Horizon Europe DMP, and DFG “Handling of Research Data”), to access a practical guide or to load a custom user-defined template. The *Import* button provides access to functions such as importing answers, clearing the user input, generating word clouds, updating front page images or loading answers from cache. The *Export* button offers options to copy the text output, export the answers to JSON, print the document to pdf or docx, update reminders, and save answers to cache. In the *Help* dropdown menu, users can access *Tutorial*, *Wiki*, *GitHub*, *Print Questions* and *Changelogs*. When users access DataPLAN for the first time, they will have the opportunity to take a step-by-step guided tour that introduces DataPLAN’s layout and functionality.

**Figure 2.**
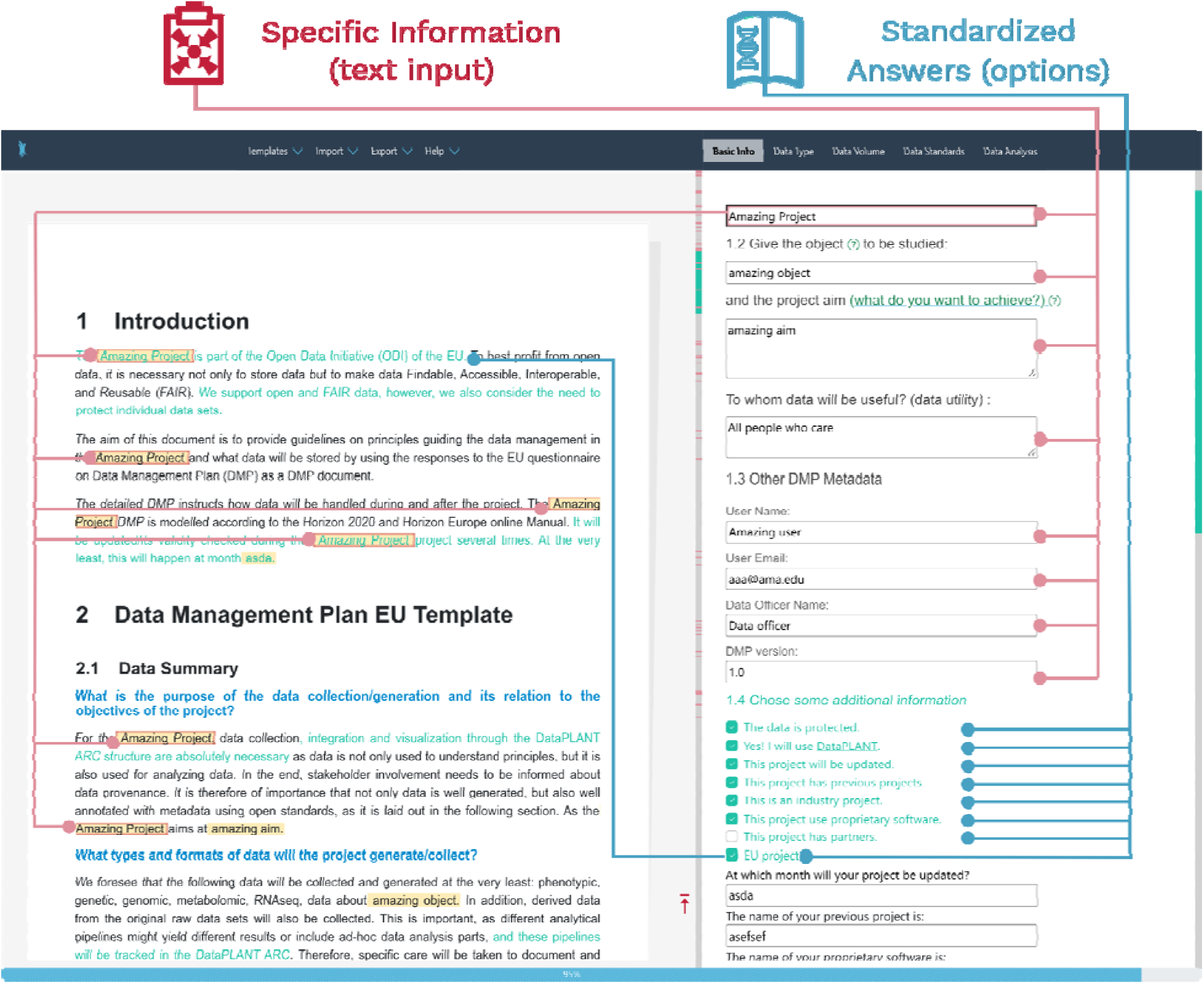
The web-based user interface of DataPLAN. The left panel displays a live preview of the DMP, while the right panel displays the DataPLAN questionnaire. In the right panel, text inputs are indicated by red lines, and blue lines represent standardized answers in checkbox format. In the left panel, red lines connect all the answers given by the first text input, while a blue line connects the answer associated with selecting the *EU project* option in the checkbox.

#### Questionnaire (right panel)

Questions can be broadly assigned to two categories: those requiring text input and those requiring the selection of one or more options from a list of checkboxes (**Figure 2, right panel**). Text input is necessary for questions about project-specific information (listed under questionnaire sections 1.1 to 1.3) such as the name of the project, the study topic, and the project’s aim, because these details vary between projects and cannot be answered using predefined options. Conversely, the questions listed under section 1.4 can be answered by selecting checkboxes, which will insert pre-determined answers into the DMP. In some instances, checkboxes may be accompanied by a text input field. For example, if the option “*This project will be updated*” is chosen in section 1.4, a prompt will appear in the questionnaire asking the user to specify the month in which the update is planned.

The SPA design, which also has no pagination in the questionnaire, is informed by the clustering analysis of the questions, which reveals partial clear clustering. Another distinctive feature is the button-less design, which enhances the user experience. The sequence of questions reflects both the order in which they appear in the DMP document and the logical progression of the research data lifecycle. For example, the *study topic* and *project aim* questions precede the *data type* question because the first two are determined earlier in the process than the latter.

#### Live preview (left panel)

The UI’s left panel displays the live preview of the DMP document (**Figure 2, left panel**). Project-specific information entered by the user is highlighted with a light yellow background, while pre-determined answers based on selected checkboxes, are distinguished by green text. Both types of text are interactive and can be hovered over or clicked. Automatic scrolling of the questionnaire is activated only when clicking the corresponding text, thus increasing convenience and minimizing the risk of errors. Additionally, when the user clicks on an answer or a question, all instances of the text are marked in red on the scroll bar, helping the user to locate the relevant information. This highlighting feature provides users with a direct visual cue, helping them to understand the relationship between the questionnaire and the resulting output.

#### DataPLAN workflow

As discussed above, DataPLAN features five steps: input, DMP generation, template change, warning/reminder, and output. Two of these (DMP generation and warning/reminder) run in the background while the others (1) provide input by answering questions (**Figure 3**, green), (2) select or customize a template (**Figure 3**, blue), and (3) collect the output (**Figure 3**, black).

**Figure 3.**
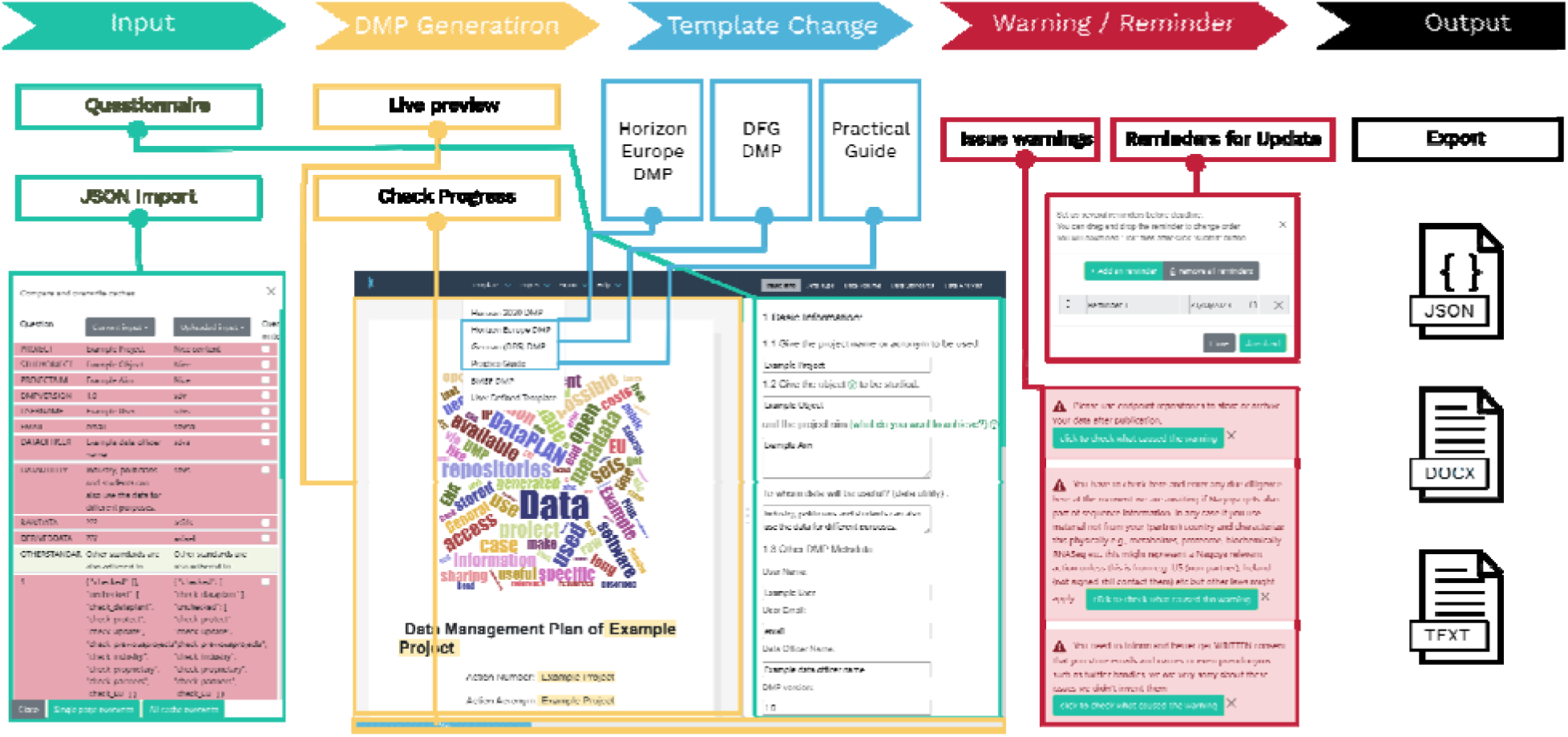
DataPLAN has five main steps. **Input (green):** Users can provide input either by completing the questionnaire manually or by importing saved input. **DMP Generation (yellow):** DMP generation (core function) loads the user input into a prepared template. **Template Change (blue):** Users can change templates at any time. We have provided users with features to help create user-defined templates. **Warning/Reminder (red):** We use warnings to make users aware of potential hurdles. The text locations that cause the warning are shown in the live preview. Reminders are downloadable ics files that can be imported into a calendar. **Output (black)**: The output of DataPLAN can be a text, docx, or JSON file.

#### Templates

The questionnaire provided by the DFG is officially described as a “checklist” [11] whereas that provided by Horizon Europe is officially described as a “template” [10]. However, the “questionnaire” is a more accurate and consistent description, so we use the terms “DFG questionnaire” and “Horizon Europe DMP questionnaire” here after. The term “template” is reserved for the DMP templates made available in DataPLAN. DataPLAN provides “Handling of Research Data” templates for DFG [11] as well as DMP templates for Horizon 2020 and Horizon Europe [10]. Because all templates can be updated simultaneously, users can select any of the templates at any point before, during, or after the input generation process (**Figure 3**, yellow). Additionally, users can customize their own template by selecting *User-defined Templates* from the *Templates* menu (**Figure 3**, blue). This allows users to modify a pre-written template in real-time to suit their needs. The modified template can then be loaded directly into the main interface and used in the same way as the others.

#### Saving and Importing data

Users can provide input either manually (by completing the questionnaire) or by importing previously saved responses. The *Export* dropdown menu features a *Download to JSON* function, allowing users to export responses in a machine- and human-readable JSON format compatible with other DMP JSON standards [52]. This enables users to import their data and resume the DMP generation process (**Figure 3**, yellow). When a JSON file is imported, the tool compares it with the current version of the DMP and presents several replacement options, such as replacing a single answer, replacing a group of cached answers, or fully replacing both the displayed and cached information.

#### Main output (DMP-related documents)

DataPLAN’s *Export* dropdown menu offers a range of options for DMP management: (1) copying text for further editing in a text editor, (2) exporting answers as a JSON file for re-use, (3) printing the document as a pdf or docx, or directly to a printer, (4) setting an update reminder by generating an ics file that can be loaded into a calendar, and (5) saving currently displayed answers to the browser cache with five available slots.

Option (1) enables users to copy one of three predefined DMP templates that fulfill the requirements of Horizon 2020, Horizon Europe, and the DFG. The text can be pasted into any text editor or printed either as a pdf or directly to a printer using option (3).

In addition to the main DMP documents, DataPLAN generates a practical guide based on user input. This guide offers detailed and optional measures to facilitate personnel task assignments, tool selection, data format conversion, and timeline arrangements. As a supplement to the main DMP, the practical guide is dynamically updated based on the user’s responses to questions. Examples of DMP documents generated by DataPLAN can be found in the **Supplementary Document S1-S4**.

#### Warnings

DataPLAN takes into account regulatory and legal considerations that arise during the creation of a DMP. For example, certain data types, such as international genomic resources and personnel data, are subject to specific regulations and laws such as the Nagoya protocol and General Data Protection Regulation (GDPR) [53,54]. To assist users navigating these complexities, DataPLAN incorporates a warning system (**Figure 3**, red) that is activated when a selected data type or format falls within the purview of these regulations. Before printing or copying the DMP, these warnings serve as timely reminders, highlighting potential challenges in data management and publication. Moreover, they include convenient links to relevant sections in the text, allowing users to access the information conveniently.

### Semantic analysis and manual comparison to standardize the content of DMPs

To identify and group similar questions in the Horizon Europe and DFG questionnaires, we conducted a semantic text similarity (STS) comparison between the two questionnaires. The questions in the two documents showed similarities ranging from 10% to 75% (**Figure 4a**). The results are provided in **Supplementary Data S1**, where each DFG question is paired with the two highest scoring questions in the Horizon Europe questionnaire, **Figure 4a** shows the complete similarity matrix as a heat map, with yellow indicating higher similarity and blue indicating lower similarity between the two questions. **Figure 4b** shows a line plot connecting DFG questions with the highest scoring Horizon Europe questions and *vice versa*, where the line color represents the similarity score (yellow color represents a higher score than blue).

**Figure 4.**
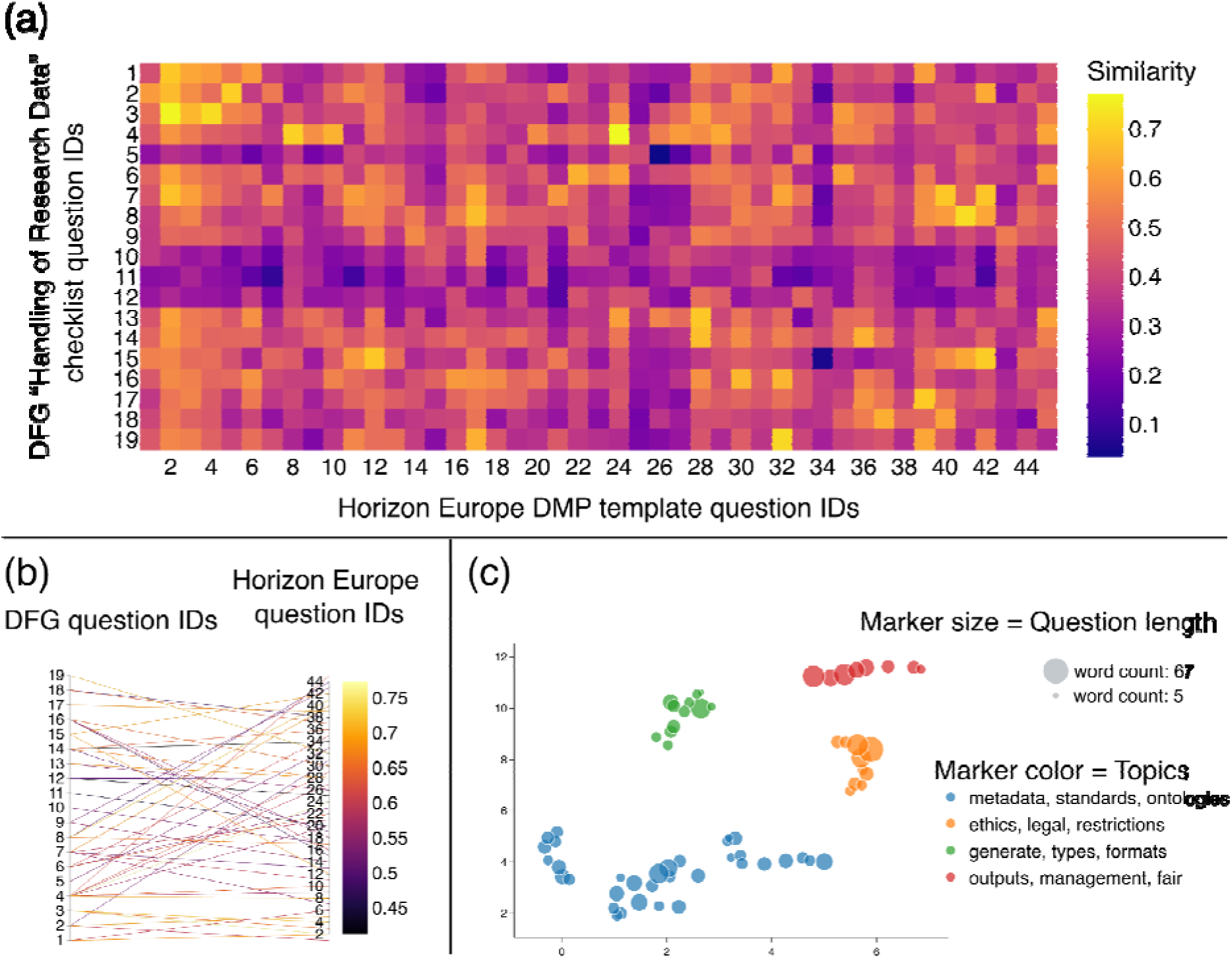
Semantic textual similarity (STS) comparison of the DFG and Horizon Europe questionnaires; (a) A heat map showed similarities between each question in the DFG and Horizon Europe questionnaires. The question IDs of the two questionnaires are shown on the two axes. The similarity between the questions is indicated with colored squares, where yellow indicates a higher similarity; (b) A parallel coordinates plot linked the two most similar questions in the DFG and Horizon Europe DMP questionnaires. The left vertical axis shows the DFG question ID and the right vertical axis shows the Horizon Europe question ID. Yellow indicates a higher similarity; (c) UMAP clustering grouped all questions in the DFG and Horizon Europe questionnaires. Four different groups of clustered scatter points showed similarities between the questions provided by each body. The axes are labeled arbitrarily, because they are outcomes of the dimension reduction process in UMAP. The radii of the scatter points represent the word count of the question. The key shows the topics of the four color-coded groups.

UMAP clustering analysis showed how the questions are related to each other (**Figure 4c**). The algorithm was able to detect close relationships between questions, with small between-cluster variance indicating similar clusters. The Calinski-Harabasz score [55] was 124.89, which is relatively low, indicating that the clusters were not well separated, and may not be distinct. There was a variation within the Calinski-Harabasz score when the assigned number of neighbors (n-neighbors) was between 2 and 19. Ultimately, we found that a value of 3 led to the highest Calinski-Harabasz score of 124.89.

The DataPLAN questionnaire design was based on the STS analysis. In our test, the Sentence-BERT model outperformed other models (**Supplementary Data S1**). Based on manual curation (**Supplementary Data S1**) we then developed the corpus for the questions and layout in DataPLAN. The different input fields for the user were designed and linked with placeholders for the three current templates.

### Testing and validating DataPLAN under FAIR principles

Testing and validating DataPLAN under FAIR principles ensures that the tool can support data management in alignment with current best RDM practices.

#### Findability

DataPLAN has implemented several measures to increase data findability. By hosting the tool on GitHub [56], developers can easily track changes in the source code and collaborate on further development. Furthermore, the use of HTML meta-tags ensures the search engine optimization of the DataPLAN website, increasing the likelihood that users will discover the tool via search engine queries. Notably, DataPLAN was included in the online registry RDMkit [57] and bio.tools [58], further augmenting its findability within the research community.

#### Accessibility

The accessibility of DataPLAN has been optimized by ensuring that it can be accessed using communications protocols that are open, free, and universally implementable. Users can access DataPLAN from desktop and mobile devices using a wide range of modern browsers, including Chrome, Firefox, Edge, Safari, and Opera. For offline work, users can download the HTML, CSS or JavaScript code using the browser’s *save* function and store them locally on their device. In addition, DataPLAN is hosted on GitHub, granting users the freedom to fork the repository and host their own personalized version. This flexibility enables users to customize and tailor the tool to suit their specific requirements. With no installation or login required, DataPLAN can be used as soon as the HTML file is downloaded, or when the website is accessed. The tool’s lightweight design and fast loading also contribute to its accessibility.

#### Interoperability

DataPLAN has been developed to ensure a high level of interoperability by enabling information exchange via JSON files, which can be read by humans and machines. The tool’s adoption of a formal, accessible, shared, and broadly applicable language for knowledge representation, along with its utilization of vocabularies that adhere to FAIR principles, further strengthens its interoperability. Additionally, DataPLAN’s purely client-side code enables easy integration into automated workflows. The ability to accept user-defined templates allows researchers to customize and tailor DMPs to their specific needs and requirements, increasing the flexibility and utility of the tool.

#### Reusability

DataPLAN’s open-source licenses allow cost-free reuse by individuals and projects [59]. The code for DataPLAN is modular, allowing developers from other fields to reuse parts of the code for their own purposes. Even people without programming skills can effortlessly create their own templates and use them to prepare DMPs. Moreover, DataPLAN’s architecture does not rely on server-side code, resulting in a seamless and easily accessible user experience. In accordance with FAIR principles, DataPLAN’s data/metadata have a detailed and transparent provenance, they are accompanied by a license that allows reuse, and they meet domain-relevant community standards, further enhancing reusability.

## Discussion

DataPLAN is a versatile but light-weight tool that facilitates the secure and efficient preparation of DMPs. DataPLAN comprises a serverless single-page web application and three DMP templates that encompass RMD best practices in the plant sciences. The templates, which incorporate 12 data types, nine endpoint repositories, and one RDM platform [24], fulfill the DMP requirements of Horizon 2020, Horizon Europe, and the “Handling of Research Data” of DFG.

### Comparison with existing DMP tools

The DMP community acknowledges the importance of maintaining a diverse range of tools [1] to meet the specific needs of different research domains. Several tools have been developed to assist users in the preparation of high-quality DMPs, such as Data Stewardship Wizard [27], DMP online [60], DMP tools [2], DMP Canvas Generator, EzDMP [37], DMPRoadmap [61], RDMO [38,42], Research Data Manager (UQRDM)[31], DataWiz [32], ARGOS [33], UWADMP [36], DMPTY [35] and easyDMP [34]. **Table 1** summarizes their key characteristics, programming languages, current templates, customizability, openness, and convenience, in comparison to DataPLAN.

**Table 1.**
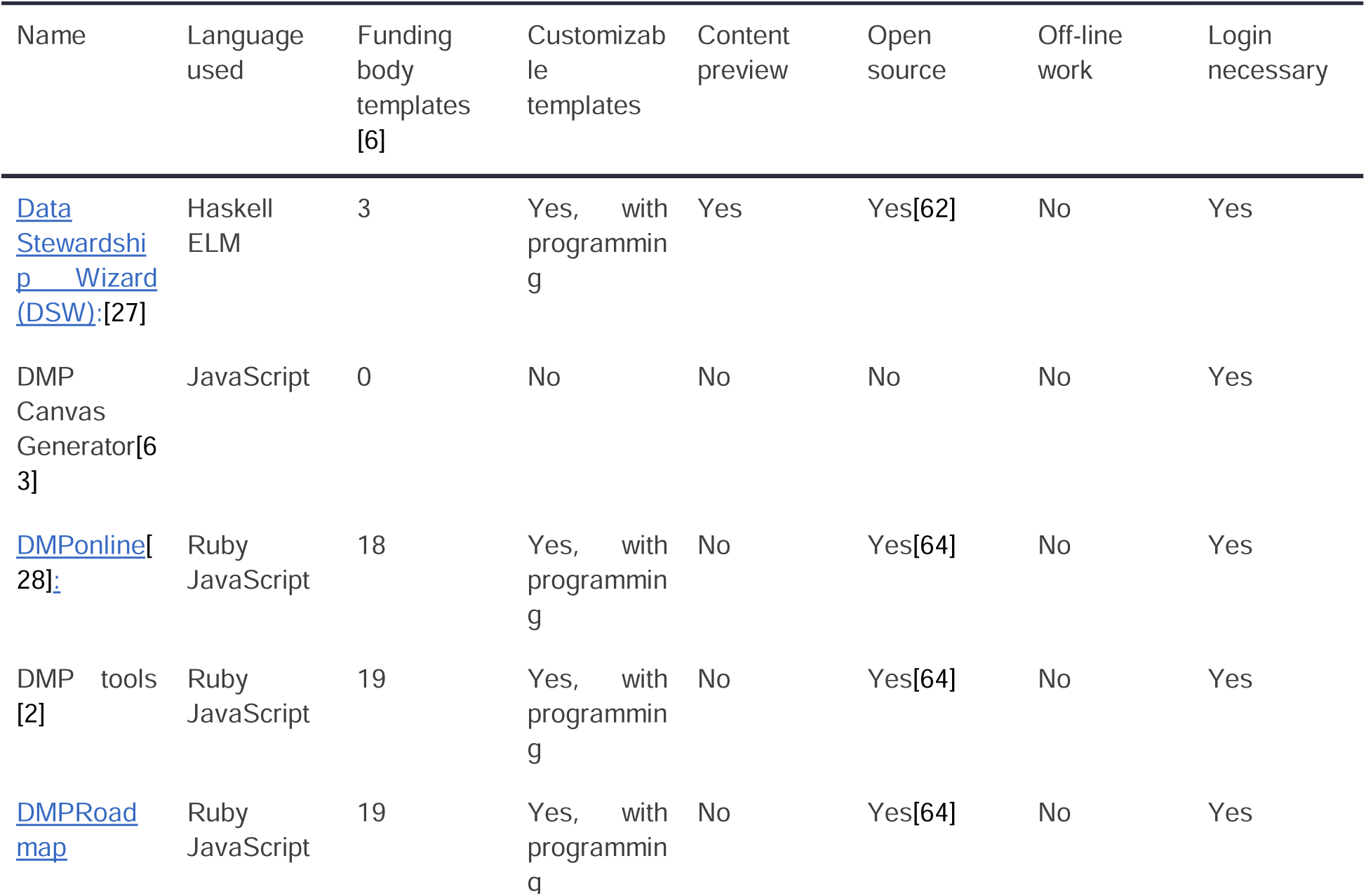
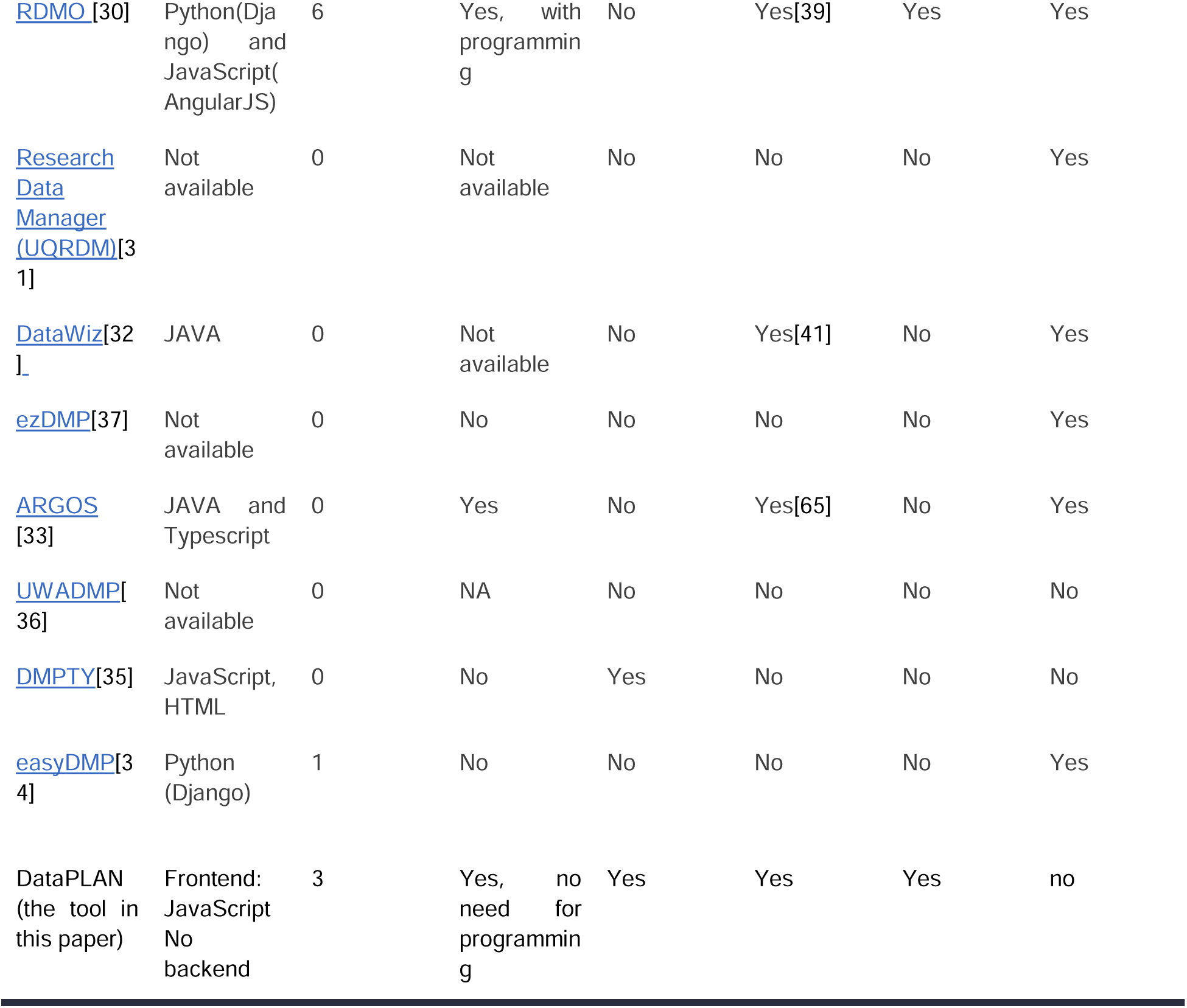
Existing DMP preparation tools compared to DataPLAN.

| Name | Language used | Funding body templates [6] | Customizable templates | Content preview | Open source | Off-line work | Login necessary |
| --- | --- | --- | --- | --- | --- | --- | --- |
| Data Stewardship Wizard (DSW): [27] | Haskell ELM | 3 | Yes, with programming | Yes | Yes[62] | No | Yes |
| DMP Canvas Generator[63] | JavaScript | 0 | No | No | No | No | Yes |
| DMPonline[28]: | Ruby JavaScript | 18 | Yes, with programming | No | Yes[64] | No | Yes |
| DMP tools [2] | Ruby JavaScript | 19 | Yes, with programming | No | Yes[64] | No | Yes |
| DMPRoadmap | Ruby JavaScript | 19 | Yes, with programming | No | Yes[64] | No | Yes |
| RDMO [30] | Python(Django) and JavaScript(AngularJS) | 6 | Yes, with programming | No | Yes[39] | Yes | Yes |
| Research Data Manager (UQRDM)[31] | Not available | 0 | Not available | No | No | No | Yes |
| DataWiz[32] | JAVA | 0 | Not available | No | Yes[41] | No | Yes |
| ezDMP[37] | Not available | 0 | No | No | No | No | Yes |
| ARGOS [33] | JAVA and Typescript | 0 | Yes | No | Yes[65] | No | Yes |
| UWADMP[36] | Not available | 0 | NA | No | No | No | No |
| DMPTY[35] | JavaScript, HTML | 0 | No | Yes | No | No | No |
| easyDMP[34] | Python (Django) | 1 | No | No | No | No | Yes |
| DataPLAN (the tool in this paper) | Frontend: JavaScript<br>No backend | 3 | Yes, no need for programming | Yes | Yes | Yes | no |

### Technical comparison

All the tools listed in **Table 1** are web-based and use JavaScript for the frontend. However, the backend programming languages vary. DataPLAN stands out because it does not require a backend at all, whereas the other tools use languages such as Python, Java, or Ruby. The pure client-side (frontend) design, which allows access in the absence of other tools or even an internet connection, is secure, because no files are transferred, making DataPLAN more reliable and trustworthy than the other tools. Furthermore, DataPLAN uses a SPA design and therefore loads a single web document and updates its content, allowing for efficient content presentation.

DataPLAN and ARGOS both provide customizable templates without coding, whereas tools like Data Stewardship Wizard, DMP online, DMP tools, DMPRoadmap, and RDMO require code to be written for customized templates. Data Stewardship Wizard, DMPTY, and DataPLAN provide a content preview, so users can see the output in real time before finishing the questionnaire. Tools such as RDMO, Data Stewardship Wizard, DMP online, DataWiz, and DataPLAN can also be used off-line. Among them, RDMO, Data Stewardship Wizard, DMP online and DataWiz require a local server, whereas DataPLAN can be used offline as soon as the website is loaded. DataPLAN, along with UWADMP, provides DMP services without needing users to log in, whereas the other tools require user registration and login.

DataPLAN thus offers a unique combination of features, including the availability of templates from multiple funding bodies, customizable templates, content preview, offline work, and no mandatory login requirement. These features enhance the usability and flexibility of DataPLAN, thus meeting the diverse needs of researchers and supporting high-quality DMP generation.

### Questionnaire and template comparison

All the tools use open-end questions (text) similar to the questionnaires provided by the funding bodies, but some tools, such as ezDMP, Data Stewardship Wizard, Open DMP, UWA DMP, RDMO and DataWiz also ask closed-end (checkbox) questions. In contrast, DataPLAN includes six closed-end questions and 10–13 open-end questions to collect project-specific information. DataPLAN offers a general questionnaire that is mapped to the templates provided by different funding bodies. Whereas other DMP tools have individual entry masks for each template, DataPLAN maintains a consistent user interface. Internally, the answers are organized and structured differently according to the selected template. This provides a streamlined user experience, eliminating the need for users to navigate different entry masks for each template.

DataPLAN goes beyond providing a blank canvas for users to fill in their DMP responses. It offers standardized pre-written answers for every question in the template (**Figure 2**, left panel). These pre-written answers are tailored to align with the specific expectations and semantic meaning of each funding body (**Figure 5**). DataPLAN provides users with the flexibility to customize these pre-written answers to meet their project’s needs (**Figure 5**), saving time and ensuring compliance with funding body guidelines. The answers integrated into the template can help users to save time, but the incorporation of a new template into DataPLAN is more time-consuming. The new template’s questions must be aligned with the established question and answer corpus within DataPLAN. Then, possible answers must be generated and included to enable integration into the DataPLAN template system. Other DMP tools [2,27,28,30] do not offer standardized answers because they must be applicable to a wide range of research domains, which can make it difficult to provide specific and practical RDM solutions. By focusing on the plant sciences, DataPLAN can provide more specific and practical answers to the questionnaire, helping researchers to create tailored DMPs. Furthermore, although the current DataPLAN template focuses on plant research, the tool can be extended to other research domains in the future.

**Figure 5.**
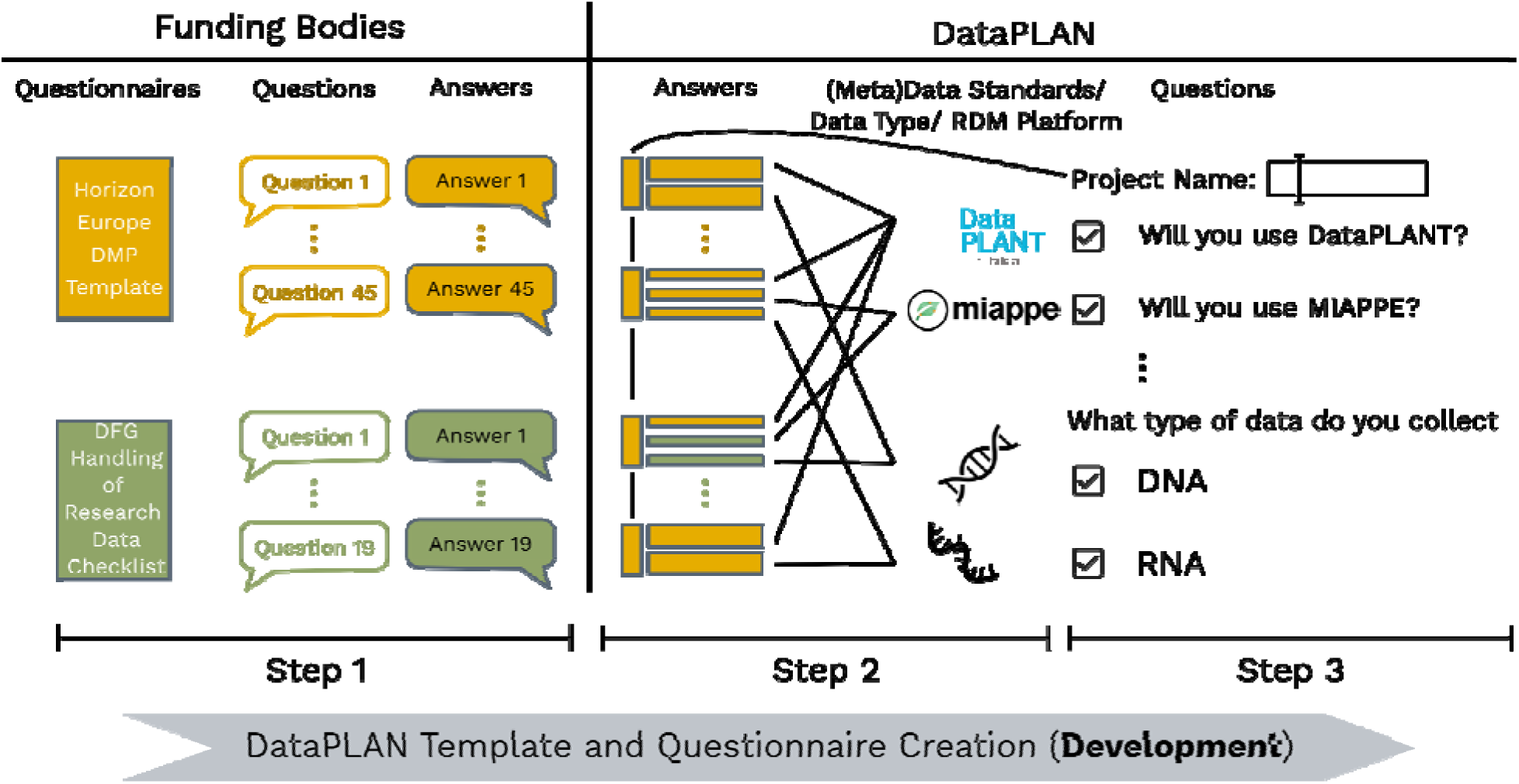
Workflow of the DataPLAN template and questionnaire design. **Step 1**: Manual checking and answering of questions in the DFG and Horizon Europe questionnaires. **Step 2**: Generation of standardized and standardized answers to fulfill the funding bodies’ requirements and align with existing metadata standards, data types and RDM platforms. **Step 3**: Design of questions that are displayed within DataPLAN.

The Data Steward Wizard offers a FAIRness analysis of the DMP by assessing the FAIRness of all user input, whereas DataPLAN provides RDM tools and platforms to improve FAIRness. RDMO, Data Steward Wizard and DataPLAN can also export DMP results in csv, xml, and/or JSON format. All the European tools (Data Stewardship Wizard, DMP online, DMP tools, DMP Canvas Generator, DMPRoadmap, RDMO, DataWiz, ARGOS, DMPTY, easyDMP, and DataPLAN) mention international agreements and regulations in their DMPs. DataPLAN includes warning notifications to make users aware of missing answers or problematic statements (e.g. those in conflict with the GDPR). It is important to be mindful of issues related to sensitive or personal information when creating a DMP, because these can have significant legal and ethical implications. If users plan to collect or store personal information as part of their research, they must obtain written informed consent from individuals and must comply with relevant laws. Failing to do so could hinder the project. Another specific feature of DataPLAN is its handling of the Nagoya Protocol [66,67], an international agreement that defines specific requirements for the sharing of and access to research data, particularly regarding genetic resources. Researchers working on projects that involve materials from developing countries that have signed the Nagoya Protocol must understand and comply with its requirements to avoid potential violations. DataPLAN assists users by providing notifications to help address legal issues and ensure research is conducted ethically and in compliance with relevant laws and regulations.

The questionnaire, template, and warnings set DataPLAN apart from other DMP tools, offering a user-friendly interface and tailored pre-written answers that enhance the efficiency and quality of the resulting DMPs.

### Integration with RDM practices

A DMP is not only a RDM practice but also encompasses planning for all other RDM tasks, fostering a higher level of interconnectedness among them [9]. For example, DMP questionnaires from funding bodies prompt users to consider data sharing even before data collection begins. This holistic approach reminds users to avoid using unsuitable tools (proprietary or non-FAIR) during data collection. The DataPLAN template facilitates this process by including RDM tools, data/metadata standards, RDM platforms and endpoint repositories (**Figure 5**, Step 2). By choosing DataPLANT [50,68] and its concepts, tools, and services such as the Annotated Research Context (ARC) [69], Swate [70], ARC Commander [71], and DataHUB [72] as the RDM practices of choice (**Figure 2**, checkbox option *Yes, I will use DataPLANT*), the resulting DMP is adapted accordingly. Consequently, if users follow the RDM practices offered by DataPLANT, this increases the FAIRness of their data.

### Integration with metadata standards

DataPLAN aligns with widely recognized minimal information standards, such as MIAME (Brazma et al., 2001), MinSEQe (Genomics Data Society, accessed on 25 June 2023), MIAPE (Taylor 2006), MSI [73] and MIAPPE [20]. These standards provide a common framework for the description and organization of data elements related to experimental planning, sample handling, and data collection/analysis. By recommending these standards, DataPLAN enables researchers to produce structured metadata, enhancing data discoverability, reusability, and reproducibility. Furthermore, the recommendations are also based on the data type selected within the DMP. Ontologies also play an important role in metadata annotations and data transformation. To enhance the annotation and transformation of metadata, DataPLAN will integrate DataPLANT Biology Ontology [74] and other relevant ontologies, as a part of the DataPLANT data management platform.

### Integration with RDM platforms

DataPLAN is designed to integrate seamlessly with RDM platforms to enhance data sharing, interoperability, and long-term preservation. The integration of DataPLAN with existing RDM infrastructures enables researchers to connect their DMPs with other RDM platforms. The integration of DataPLANT [50,68] and its tools and concepts (such as ARC, Swate [70], ARC Commander [71], and DataHUB [72]) into DataPLAN templates can streamline and simplify data management for researchers. The tools and resources cover every stage of RDM, from data acquisition to publication. By providing links and guides for the use of such resources within DataPLAN, researchers can access the tools and resources more easily, allowing them to manage their data more effectively. DataPLAN also provides a step-by-step guide, helping researchers to use RDM platforms for data management throughout their research projects. The integration of RDM platforms into DataPLAN enhances the efficiency and effectiveness of data management for researchers.

### Outlook

DataPLAN currently meets the requirements of H2020, Horizon Europe, and DFG projects in the plant sciences. However, international interdisciplinary projects may have different preferences and may require a cloud-based collaborative environment for DMP preparation in the future. DataPLAN will therefore be made more compatible with other DMP tools and will incorporate additional domain specific templates. We will also continuously integrate new resources such as tools, ontologies, and infrastructures.

### Integration with DMP and RDM tools

Among the 13 tools listed in **Table 1**, the input and output of RDMO will be supported by DataPLAN in the future. Templates in RDMO and Data Stewardship Wizard will be also very helpful for future template development. Domain specific templates such as biodiversity and emission RDMO templates [75,76] are helpful resources that will be referenced for additional template development. By ensuring compatibility with other tools and considering the DMP needs of other domains, the DataPLAN will gain more users and become more helpful for the community.

Software catalogs such as FAIRsharing [77] and bio.tools [58] have been developed and maintained to provide a list of RDM tools. DataPLAN will use bio.tools and FAIRsharing to propose appropriate tools based on the scientific field and data type specified within the DMP. Such suggestions during the RDM planning phase could prevent issues caused by tool incompatibilities. Moreover, by selecting appropriate tools at an earlier stage, it will be possible to incorporate more intricate workflows and pipelines into the DMP.

### Integration with data analysis tools

To streamline data management, DataPLAN will enhance its integration with popular data analysis tools and platforms. This will enable researchers to seamlessly connect their DMPs with data analysis workflows, data visualization tools, and statistical analysis software. By linking DMPs with newly developed data analysis tools, researchers can ensure that their RDM practices align with the most advanced data analysis tools and processing requirements.

### Collaboration, version control and institutional integration

DataPLAN will enhance its collaboration features, allowing researchers to collaborate on DMP development and maintenance without using GitHub or GitLab. This includes functionalities such as version control, comments, and sharing capabilities. These collaborative features will enable research teams to collectively develop, review, and refine DMPs, promoting better RDM practices and fostering collaboration between research projects.

DataPLAN plans to integrate with institutional repositories to facilitate the seamless deposition and publication of data. This integration will enable researchers to directly link their DMPs with the corresponding datasets in institutional repositories. By automating the process, researchers can ensure that the datasets associated with their DMPs are readily available for sharing and discovery, fostering open and transparent research practices.

## Methods

### Software development

The DataPLAN software was implemented in client-side (frontend) JavaScript [78] as a standalone single page application (SPA), which can be executed in all modern web browsers (e.g. Google Chrome, Microsoft Edge, Mozilla Firefox, Opera and Safari). As a client-side SPA, data security is ensured, because no software installation is required and no information is sent to external servers. For implementation, several open source libraries are used in the user interface (UI) and in visualizations: Bootstraps 5 [79], bs5-intro-tour [80], d3 word cloud [81], FileSaver [82], and split.js [83]. DataPLAN can be accessed via the DataPLANT website (https://plan.nfdi4plants.org/; accessed on 25 June 2023).

*The DMP Generation* process was the highest priority during DataPLAN development. To generate the DMP output, the entire template is searched multiple times. Following each search, a portion of the text is modified based on the user input. Partial text modification is achieved through the use of two distinct types of placeholders: first, a regular placeholder, denoted by the $_ symbol, which simply inserts the user’s text into the sentence; and second, the use of more complex structures, where placeholders are embedded within conditional statements (if-structures) that are controlled by either user input or the choice of template. These placeholders are identified using the window.find() function, with the search range extended to the left by three characters in order to locate the relevant conditional statement (#if or #if!). Additionally, once the #if has been found, the right boundary #endif must also be located. When these conditions have been identified, a decision tree determines the appropriate changes to be made based on the user input. The *Template Change* and *Warning/Reminder* sections were developed next, and the *Input* and *Output* sections last. The five sections can also be run and developed in parallel to ensure that the *DMP Generation* process is robust and reliable.

### DMP template and questionnaire design

We created DMP templates that fulfill the requirements of DFG and Horizon Europe in three consecutive steps. First, all questions provided by the DFG and Horizon Europe questionnaires were answered manually (**Figure 2**, Step 1). Second, the answers were adapted to align with typical data/metadata standards or data types used in the plant sciences (**Figure 2**, Step 2). Third, in order to standardize DataPLAN’s answers for different technologies, we also studied underlying data generation and analysis methods, data formats, known minimum information standards and repositories for data publishing (**Table 2**). The related information (e.g. which minimum information standard should be used to report metadata from a specific high-throughput technology) was incorporated into the templates. This enabled us to generate DMPs that are both standardized and domain-specific. All the answers were used to design templates that (a) fulfill the requirements of DFG and Horizon Europe and (b) allow the selection of optional answers based on data standards or data types (**Figure 2**).

**Table 2.**
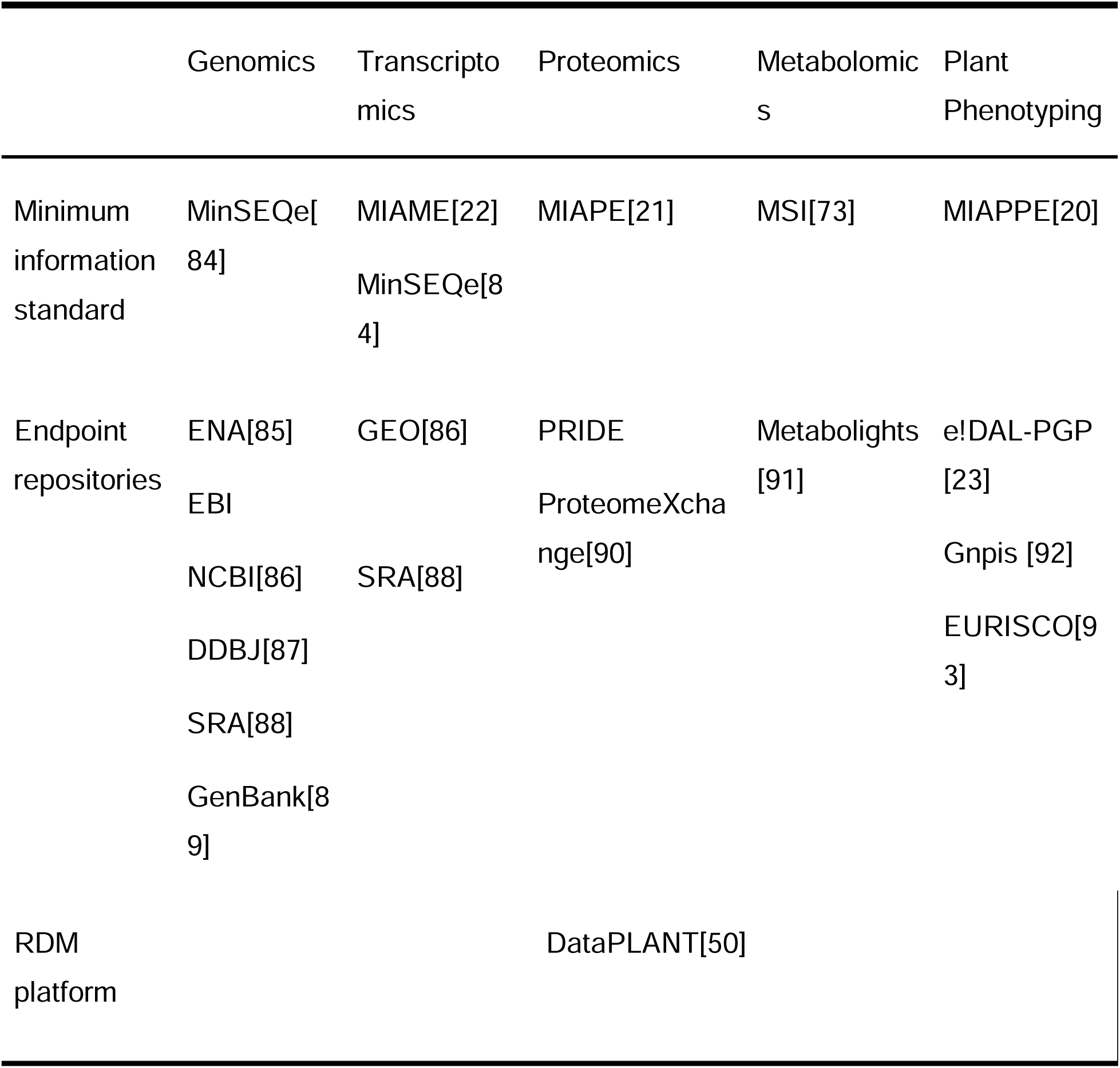
A collection of minimum information standards, endpoint repositories and data management platforms relevant to DataPLAN.

|  | Genomics | Transcriptomics | Proteomics | Metabolomics | Plant Phenotyping |
| --- | --- | --- | --- | --- | --- |
| Minimum information standard | MinSEQe[84] | MIAME[22]<br>MinSEQe[84] | MIAPE[21] | MSI[73] | MIAPPE[20] |
| Endpoint repositories | ENA[85]<br>EBI<br>NCBI[86]<br>DDBJ[87]<br>SRA[88]<br>GenBank[89] | GEO[86]<br>SRA[88] | PRIDE<br>ProteomeXchange[90] | Metabolights [91] | e!DAL-PGP [23]<br>Gnpis [92]<br>EURISCO[93] |
| RDM platform | DataPLANT[50] |  |  |  |  |

To design the questionnaire, similar questions in the questionnaires of Horizon Europe and DFG (**Figure 5**, analysis between Steps 1 and 2) were identified by semantic analysis using the pre-trained Sentence-BERT model [94]. This model computes the cosine-similarity score between each question in the DFG and Horizon Europe questionnaires. The results were manually (**Figure 5**, Step 3) verified for meaningfulness and relevance within the context of data management and the plant sciences. Clustering was then applied using the dimension reduction software UMAP [95], and the clustering software HDBSCAN [96].

### Testing

In order to ensure the proper functioning of DataPLAN, we implemented automated technical testing and user testing procedures. For automated testing, we used PerformanceMeasure [97] and the web development tool LightHouse [98]. Manual testing involved two steps: first, developer testing for core functionalities such as placeholder replacement, logic condition of placeholders, and template switch; and second, testing by potential users with experience in the plant sciences for aspects such as usability, time expenditure, user interface, template selection, customizability, RDM information and best practices such as guidelines, collaborative features, version control, compliance checking, and the import and export of DMPs.

## Conclusions

External platforms such as GitHub or GitLab are used for collaborative communication in DataPLAN. This weakness in collaboration is a disadvantage of pure frontend designs. The collaborative functions can be enhanced by adding an independent optional backend. This independent frontend (client-side) and backend (server-side) architecture makes DataPLAN unique among existing tools. We applied machine learning in the development of DMP tools, resulting in an objective and reproducible interpretation on the need for DMPs. DataPLAN is a unique, lightweight tool that emphasizes FAIR principles, and puts them into practice to meet the current DMP needs of funding bodies in the plant sciences.

We have developed DataPLAN, which enables the preparation and updating of DMPs requested by multiple funding bodies. Equipped with pre-written standardized answers and a single-page interface, DataPLAN enables researchers to create standardization DMPs in minutes, regardless of their experience and expertise. We use a pure frontend design to prevent data transmission, enhancing the security and privacy of the data. DataPLAN is an open-source and web-based tool, enabling customization by users and modification by developers. Overall, DataPLAN is a valuable resource for researchers seeking to manage their data efficiently while maintaining high standards of security and privacy.

## Supplementary Materials

Data S1: Semantic text similarity comparison and manual curation of questions in DFG and Horizon Europe questionnaires, Document S1: Expanded description of the tool functions and analysis, Document S2: Example DataPLAN DMP output of a H2020 project, Document: S3: Example DataPLAN DMP output of a Horizon Europe project, Document S4: Example DataPLAN DMP output of a DFG project

## Author Contributions

Conceptualization, B.U. and X.-R.Z.; methodology, X.-R.Z. and S.B.; software, X.-R.Z.; validation, X.-R.Z., S.B., D.B., C.M.R and A.K.; formal analysis, X.-R.Z. and S.B.; investigation, X.-R.Z. and S.B.; resources, B.U.; data curation, X.-R.Z., S.B., R.M.T. and D.B.; writing—original draft preparation, X.-R.Z., S.B., D.B., B.U. and A.K.; writing—review and editing, R.M.T., S.B., D.B., B.U., C.M.R., T.M., D.V.S., A.K. and X.-R.Z.,; visualization, X.-R.Z.; supervision, A.K.; project administration, A.K. and C.M.R.; funding acquisition, D.V.S., B.U., and T.M. All authors have read and agreed to the published version of the manuscript.

## Acknowledgement

We would also like to thank our colleagues at DataPLANT and IBG-4 for their feedback and support throughout the research process. In particular, we would like to thank Dr. Elisa Senger, Dr. Andrea Schrader, Dr. Kathryn Dumschott and Hannah Dörpholz for their valuable insights and suggestions.

## List of Abbreviations

ARC: Annotated Research Context
DDBJ: DNA Data Bank of Japan
DFG: German Research Foundation (Deutsche Forschungsgemeinschaft)
DMP: Data management plan
EBI: European Bioinformatics Institute
ENA: European Nucleotide Archive
EU: European Union
FAIR: Findable, accessible, interoperable and reusable
GDPR: EU General Data Protection Regulation
GEO: Gene Expression Omnibus
LLM: Large language model
MIAME: Minimal information about a microarray experiment
MIAPE: Minimum information about a proteomics experiment
MIAPPE: Minimal information about plant phenotyping experiment
MinSEQe: Minimum information about a high-throughput sequencing experiment
MSI: Metabolomics Standards Initiative
NCBI: National Center for Biotechnology Information
NFDI: National Research Data Infrastructure (of Germany)
PRIDE: Proteomics Identification Database
RDM: Research data management
RNA-Seq: RNA sequencing
SRA: Sequence Read Archive
STS: Semantic textual similarity
UI: User interface

