## Supplementary material for "DataPLAN: a web-based data management plan generator for the plant sciences": Document_S1.doc

Export HTML To Doc

Data Management Plan of the H2020 Project  Example Project

|  |  |
| --- | --- |
| Action Number: | Example Project |
| Action Acronym: | Example Project |
| Action Title: | Example Project |
| Date: | July 7, 2023 |
| DMP version: | 1.0 |

### 1    Introduction

To best profit from open data, it is necessary not only to store the data but to make it Findable, Accessible, Interoperable, and Reusable (FAIR).

The aim of this document is to provide guidelines on the principles of data management in the  Example Project  and to specify which type of data will be stored, this will be achieved by using the responses to the EU questionnaire on Data Management Plan (DMP) as a DMP
document.

The detailed DMP states how data will be handled during and after the project. The  Example Project  DMP is prepared according to the Horizon 2020 and Horizon Europe online manual.

### 2    Data Management Plan EU Template

#### 2.1    Data Summary

What is the purpose of the data collection/generation and its relation to the objectives of the project?

The  Example Project  has the following aim:  Example Aim.  Therefore, data collection, integration and visualization   through a standardized data management process is absolutely necessary  because the data are used not only to understand principles, but also be informed about the provenance of data analyzing data. Stakeholders must also be informed about the provenance of data. It is therefore necessary to ensure that the data are well generated and also well annotated with metadata using open standards, as laid out in the next section.

What types and formats of data will the project generate/collect?

The  Example Project  will collect and/or generate the following types of raw data : phenotypes, genotypes, other NGS data, metabolome, RNA-Seq and other forms of transcriptomic data, data related  Example Topic.  In addition, the raw data will also be processed and modified using analytical pipelines, which may yield different results or include ad hoc data analysis parts. . Therefore, care will be taken to document and archive these resources (including the analytical pipelines) as well.

Will you re-use any existing data and how?

The project builds on existing data sets and relies on them.  For example, without a proper genomic reference it is very difficult to analyze next-generation sequencing (NGS) data sets. It is also important to
include existing data-sets on the expression and metabolic behavior of the  Example Topic, and on existing background knowledge.
Genomic references can be gathered from reference databases for genomes/ and sequences, like the US National Center for Biotechnology Information: NCBI, European Bioinformatics Institute: EBI; DNA Data
Bank of Japan: DDBJ. Furthermore, prior 'unstructured' data in the form of publications and data contained therein will be used for decision making.

What is the origin of the data?

Public data will be extracted as described in the previous paragraph. For the  Example Project, specific data sets will be generated by the consortium partners.

Data of different types or representing different domains will be generated using unique approaches. For example:

- RNA sequencing will be generated using shor-read or long-read plantforms, either in house or outsourced to academic facilities or commercial services, and the raw data will be processed using estabilished biofirmatics piplines.
- Metabolomic data will be generated by coupled chromatography and mass spectrometry using targeted or untarget approaches.
 - Proteomic data will be generated using coupled chromatography and mass spectrometry for the analysis of protein abundance and protein identification, as well as additional techniques for structural analysis, the identification of post-translational modifications and the characterization of protein interactions.
- Image data will be generated by equipment such as cameras, scanners, and microscopes combined with software. Original images which contain metadata such as exif photo information will be archived.
- Genomic data will be created from sequencing data, which will be processed to identify genes, regulatory elements, transposable elements, and physical markers such as SNPs, microsatellites and structural variants.
- Genetic data will be generated targeting crosses and breeding experiments, and will include recombination frequencies and crossover event that position genetic markers and quantitative trait loci that can be associated with physical genomic markers/variants.
- Targeted assays data (e.g. glucose and fructose concentrations or production/ultilization rates) will be generated using specific equipment and methods that are fully documented in the laboratory notebook.
- Model data will be generated by using software simulations. The complete workflow, which includes the environment, runtime, parameters, and results, will be documented and archived.
- Computer code will be produced by programmers.
- Excel data will be generated by data analysts by using MS Office or open-source software.
- The origin and assembly of cloned DNA will include (a) source of original vector sequence with Add gene reference where available, and source of insert DNA (e.g., amplification by PCR from a given sample, or obtained from existing library), (b) cloning strategy (e.g., restriction endonuclease digests/ligation, PCR, TOPO cloning, Gibson assembly, LR recombination), and (c) verified DNA data sequence of final recombinant vector.
- Phenotypic data will be generated using phenotyping platforms and corresponding ontologies, including number/size of organs such as leaves, flowers, buds etc., size of whole plant, stem/root architecture (number of lateral branches/roots etc), organ structures/morphologies, quantitative metrics such as color, turgor, health/nutrition indicators, among others.

What is the expected size of the data?

We expect to generate  ???  GB of raw data and up to  ???  GB of processed data.

To whom might it be useful ('data utility')?

The data will initially benefit the  Example Project  partners, but will also be made available to selected stakeholders closely involved in the project, and then the scientific community working on  Example Topic.   Industry, politicians and students can also use the data for different purposes.  In addition, the general public interested in  Example Topic  can also use the data after publication. The data will be disseminated according to the  Example Project' s dissemination and communication plan,

#### 2.2    FAIR data

##### Making data findable, including provisions for metadata

Are the data produced and/or used in the project discoverable with metadata, identifiable and locatable by means of a standard identification mechanism (e.g. persistent and unique identifiers such as Digital
Object Identifiers)?

All datasets will be associated with unique identifiers and will be annotated with metadata. The  Example Project  will rely on community standards plus additional recommendations applicable in the plant science, such as the Minimum Information About a Plant Phenotyping Experiment (MIAPPE).  
Unlike cross-domain minimal sets such as the Dublin core, which mostly define the submitter and the general type of data, allow reusability by other researchers by defining properties of the plant (see the preceding section). However, minimal cross-domain annotations also remain part of the  Example Project.    Other standards are also adhered to.  

What naming conventions do you follow?

Data variables will be allocated standard names. For example, genes, proteins and metabolites will be named according to approved nomenclature and conventions. These will also be linked to functional ontologies where possible. Datasets will also be named I a meaningful way to ensure readability by humans. Plant names will include traditional names, binomials, and all strain/cultivar/subspecies/variety identifiers.

Will search keywords be provided that optimize possibilities for re-use?

Keywords about the experiment and consortium will be included, as well as an abstract about the data, where useful. In addition, certain keywords can be auto-generated from dense metadata and its
underlying ontologies.

Do you provide clear version numbers?

To maintain data integrity and facilitate reanalysis, data sets will be allocated version numbers where this is useful (e.g. raw data must not be changed and will not get a version number and is considered
immutable).

What metadata will be created? In case metadata standards do not exist in your discipline, please outline what type of metadata will be created and how.

We will use Investigation, Study, Assay (ISA) specification for metadata creation. For specific data (e.g., RNASeq or genomic data), we use metadata templates from the end-point repositories.  The Minimum Information About a Next-generation Sequencing Experiment (MinSEQe) will also be used. 

  Metabolights submission compliant standards will be used for metabolomic data where this is acccepted by the consortium partners.<--pay attention to the sentence before because some Metabolomics partners considers Metabolights
not an accepted standard As a part of plant research community, we use  MIAPPE for phenotyping data in the broadest sense, but we will also be rely on  specific SOPs for additional
annotations

##### Making data openly accessible

Which data produced and/or used in the project will be made openly available as the default? If certain datasets cannot be shared (or need to be shared under restrictions), we explain why, clearly separating
legal and contractual reasons from voluntary restrictions.

By default, all data sets from the  Example Project  will be shared with the community and made openly available. However, before the data are released, all will be provided with an opportunity to check for potential IP (according to the consortium agreement and background IP rights). IP protection will be prioritized for datasets that offer the potential for exploitation.

Note that in multi-beneficiary projects it is also possible for specific beneficiaries to keep their data closed if relevant provisions are made in the consortium agreement and are in line with the reasons
for opting out.

How will the data be made accessible (e.g. by deposition in a repository)?

Data will be made available via the  Example Project  platform using a user-friendly front end that allows data visualization. Besides this it will be ensured that data which can be stored in
international discipline related repositories which use specialized technologies (Sequencing at the national US center Sequencing and sequence data at the EU center: EBI, Proteome database:
metabolomic database: ) will be used to store data and the data will be processed there as well.

Specialized repositories will be used where appropriate, such as INSDC (GenBank, EBI, DDBJ) for nucleotide sequence data, PIR/UniProt/SWISS-PROT for proteins, PDB for protein structures, GEO for transcriptomics, PRIDE for proteomics data, and METLIN for metabolomics data. For unstructured and less standardized data (e.g., experimental phenotypic measurements), these will be annotated with metadata and if complete allocated a digital object identifier (DOI).

What methods or software tools are needed to access the data?

No specialized software will be needed to access the data, just a modern browser. Access will be possible through web interfaces. For data processing after obtaining raw data, typical
open-source software can be used.

Is documentation about the software needed to access the data included?

All external software documentation will be duplicated locally and stored near the software.

Is it possible to include the relevant software (e.g. in open-source code)?

As stated above, the  Example Project  will use publicly available open-source and well-documented certified software .

Where will the data and associated metadata, documentation and code be deposited? Preference should be given to certified repositories that support open access, where possible.

As noted above, specialized repositories will be used for common data types. For unstructured and less standardized data (e.g., experimental phenotypic measurements), these will be annotated with metadata and if complete allocated a digital object identifier (DOI)..

Have you explored appropriate arrangements with the identified repository?

The submission is for free, and it is the goal (at least of ENA) to obtain as much data as possible. Therefore, arrangements are neither necessary nor useful. Catch-all repositories are not required.
.

If there are restrictions on use, how will access be provided?

There are no restrictions beyond the IP screening described above, which is in line with European open data policies.

Is there a need for a data access committee?

There is no need for a data access committee.

Are there well described conditions for access (i.e. a machine-readable license)?

Yes, where possible, e.g. CC REL will be used for data not submitted to specialized repositories such as ENA.

How will the identity of the person accessing the data be ascertained?

Where data are shared only within the consortium, if the datasets are not yet finished or are undergoing IP checks, the data will be hosted internally and a username and password will be required for access (see GDPR rules). When the data are made public in EU or US repositories, completely anonymous access is normally allowed. This is the case for ENA as well and both are in line with GDPR requirements.

##### Making data interoperable

Are the data produced in the project interoperable, that is allowing data exchange and re-use between researchers, institutions, organizations, countries, etc. (i.e. adhering to standards for formats, as much
as possible compliant with available (open) software applications, and in particular facilitating re-combinations with different datasets from different origins)?

Whenever possible, data will be stored in common and openly defined formats including all the necessary metadata to interpret and analyze data in a biological context. By default, no proprietary formats will be
used. However Microsoft Excel files (according to ISO/IEC 29500-1:2016) might be used as intermediates by the consortium. In addition, text
files might be edited in text processor files, but will be shared as pdf.

What data and metadata vocabularies, standards or methodologies will you follow to make your data interoperable?

As noted above, we foresee using minimal standards such as   MinSEQe for sequencing data and  Metabolights compatible forms for metabolites 
and MIAPPE for phenotyping-like data . The minimal information standards will allow the integration of data across projects, and its reuse according to established and tested protocols. We will also use
ontological terms to enrich the data sets relying on free and open ontologies where possible. Additional ontology terms might be created and canonized during the  Example Project. 

Will you be using standard vocabularies for all data types present in your data set, to allow inter-disciplinary interoperability?

Open ontologies will be used where they are mature. As stated above, some ontologies and controlled vocabularies might need to be extended.

In case it is unavoidable that you use uncommon or generate project specific ontologies or vocabularies, will you provide mappings to more commonly used ontologies?

Common and open ontologies will be used, so this question does not apply.

##### Increase data reuse (by clarifying licences)

How will the data be licensed to permit the widest re-use possible?

Open licenses, such as Creative Commons (CC), will be used whenever possible.

When will the data be made available for re-use? If an embargo is sought to give time to publish or seek patents, specify why and how long this will apply, bearing in mind that research data should be made
available as soon as possible.

The data will be published as soon as possible to guarantee reusability.   IP issues will be checked before publication.  All consortium partners will be
encouraged to make data available before publication, openly and/or under pre-publication agreements  such as those started in Fort Lauderdale and set forth by the Toronto International Data
Release Workshop.  This will be implemented as soon as IP-related checks are complete.

Are the data produced and/or used in the project usable by third parties, in particular after the end of the project? If the re-use of some data is restricted, explain why.

There will be no restrictions once the data are made public.

How long is it intended that the data remains re-usable?

The data will be made available for many years.

Data submitted to repositories (as detailed above) e.g. ENA /PRIDE would be subject to local data storage regulation.

Are data quality assurance processes described?

The data will be checked and curated. .

#### 2.3    Allocation of resources

What are the costs for making data FAIR in your project?

The  Example Project  will bear the costs of data curation, and data maintenance/security before transfer to public repositories. Subsequent costs are then borne by the operators of these repositories.

Additionally, costs for after publication storage are incurred by end-point repositories (e.g. ENA) but not charged against the  Example Project  or its members but by the operation budget of these repositories.

How will these be covered? Note that costs related to open access to research data are eligible as part of the Horizon 2020 or Horizon Europe grant (if compliant with the Grant Agreement conditions).

The cost born by the  Example Project  are covered by the project funding. Pre-existing structures will also be used.

Who will be responsible for data management in your project?

The responsible person will be  Example data officer name  of the  Example Project. 

Are the resources for long term preservation discussed (costs and potential value, who decides and how/what data will be kept and for how long)?

The data officer will ultimately decides on the strategy to preserve data that are not submitted to end-point subject area repositories when the
project ends. This will be in line with EU guidlines, institute policies, and data sharing based on EU and international standards.

#### 2.4    Data security

What provisions are in place for data security (including data recovery as well as secure storage and transfer of sensitive data)?

Online platforms will be protected by vulnerability scanning, two-factor authorization and daily automatic backups allowing immediate recovery. All partners holding confidential project data to use secure platforms with automatic backups and offsite secure copies.

Is the data safely stored in certified repositories for long term preservation and curation?

Wherever there are certified repositories, these will be used as end-point repositories.  Transcriptomics data and gene sequence data will be also made available upon publication via the standards
ENA/SRA,   metabolite data in e.g. Metabolights (and/or Nationwide repositories like the German NFDI the French INRAe),   Proteomics data in
e.g. Pride/Proteomexchange . In addition, the national resource will maintain safekeeping of data also after the project ends.

#### 2.5    Ethical aspects

Are there any ethical or legal issues that can have an impact on data sharing? These can also be discussed in the context of an ethics review. If relevant, include references to ethics deliverables and
ethics chapter in the Description of the Action (DoA).

At the moment, we do not anticipate ethical or legal issues with data sharing. In terms of ethics, since this is plant data, there is no need for an ethics committee to deal with data from plants, although we will diligently follow the Nagoya protocol on access and benefit sharing. (🡺see Nagoya protocol). <--pay attention to the sentence before because You have to check here and enter any due diligence here at the moment we are awaiting if Nagoya gets also part of sequence information. In any case
if you use material not from your (partner) country and characterize this physically e.g., metabolites, proteome, biochemically RNASeq etc. this might represent a Nagoya relevant action unless this is from e.g.
US (non partner), Ireland (not signed still contact them) etc but other laws might apply….

Is informed consent for data sharing and long term preservation included in questionnaires dealing with personal data?

The only personal data that will potentially be stored is the submitter name and affiliation in the metadata for data. In addition, personal data will be collected for dissemination and communication
activities using specific methods and procedures developed by the  Example Project  partners to adhere to data protection. <--pay attention to the sentence before because You need to inform and better get WRITTEN consent that you store emails and
names or even pseudonyms such as twitter handles, we are very sorry about these issues we didn’t invent them

#### 2.6    Other issues

Do you make use of other national/funder/sectorial/departmental procedures for data management? If yes, which ones?

Yes, the  Example Project  will use common Research Data Management (RDM) tools.

### 3     Annexes

#### 3.1     Abbreviations

CC Creative Commons

CC CEL Creative Commons Rights Expression Language

DDBJ DNA Data Bank of Japan

DMP Data Management Plan

DoA Description of Action

DOI Digital Object Identifier

EBI European Bioinformatics Institute

ENA European Nucleotide Archive

EU European Union

FAIR Findable Accessible Interoperable Reproducible

GDPR General data protection regulation (of the EU)

IP Intellectual Property

ISO International Organization for Standardization

MIAMET Minimal Information about Metabolite experiment

MIAPPE Minimal Information about Plant Phenotyping Experiment

MinSEQe Minimum Information about a high-throughput Sequencing Experiment

NCBI National Center for Biotechnology Information

NFDI National Research Data Infrastructure (of Germany)

NGS Next Generation Sequencing

RDM Research Data Management

RNASeq RNA Sequencing

SOP Standard Operating Procedures

SRA Short Read Archive

ONP Oxford Nanopore

qRTPCR quantitative real time polymerase chain reaction

WP Work Package
