## Supplementary material for "DataPLAN: a web-based data management plan generator for the plant sciences": Document_S2.doc

Export HTML To Doc

Data Management Plan of the Horizon Europe Project  Example Project

|  |  |
| --- | --- |
| Action Number: | Example Project |
| Action Acronym: | Example Project |
| Action Title: | Example Project |
| Date: | July 7, 2023 |
| DMP version: | 1.0 |

### Introduction

To best profit from open data, it is necessary not only to store data but to make data Findable, Accessible, Interoperable, and Reusable (FAIR).

The aim of this document is to provide guidelines on principles guiding the data management in the  Example Project  and what data will be stored by using the responses to the EU questionnaire on Data Management Plan (DMP) as a DMP
document.

The detailed DMP instructs how data will be handled during and after the project. The  Example Project  DMP is modified according to the Horizon Europe and Horizon Europe online Manual.

#### 1.    Data Summary

Will you re-use any existing data and what will you re-use it for? State the reasons if re-use of any existing data has been considered but discarded.

The project builds on existing data sets and relies on them.  For instance, without a proper genomic reference it is very difficult to analyze NGS data sets. It is also important to include existing data sets on the expression and metabolic behaviour of  Example Topic, but of course, also on existing characterization and the background knowledge. Genomic references can simply be gathered from reference databases for genomes/sequences, like the National Center for Biotechnology Information: NCBI (US); European Bioinformatics Institute: EBI (EU); DNA Data Bank of Japan: DDBJ (JP). Furthermore, prior 'unstructured' data in the form of publications and data contained therein will be used for decision making.

What types and formats of data will the project generate or re-use?

The  Example Project  will collect and/or generate the following types of raw data: phenotypes, genotypes, other NGS data, metabolome, RNA-Seq and other forms of transcriptomic data, data about  Example Topic.  In addition, derived data from the
original raw data sets will also be collected. This is important, as different analytical pipelines might yield different results or include ad-hoc data analysis parts,. Therefore, specific care will be taken to document and archive these resources (including the analytic pipelines) as well. 

What is the purpose of the data generation or re-use and its relation to the objectives of the project?

The  Example Project  has the following aim:  Example Aim.  Therefore, data collection, integration and visualization   through a standardized data management process is absolutely necessary  because the data are used not only to understand principles, but also be informed about the provenance of data analyzing data. Stakeholders must also be informed about the provenance of data. It is therefore necessary to ensure that the data are well generated and also well annotated with metadata using open standards, as laid out in the next section.

What is the expected size of the data that you intend to generate or re-use?

We expect to generate raw data in the range of  ???  GB of data. The size of the derived data will be about  ???  GB.

What is the origin/provenance of the data, either generated or re-used?

  Industry, politicians and students can also use the data for different purposes.
 

#### 2    FAIR data

##### 2.1. Making data findable, including provisions for metadata

Will data be identified by a persistent identifier?

All data sets will receive unique identifiers, and they will be annotated with metadata.

Will rich metadata be provided to allow discovery? What metadata will be created? What disciplinary or general standards will be followed? In case metadata standards do not exist in your discipline, please outline what type of metadata will be created and how.

The  Example Project  will rely on community standards plus additional recommendations necessary in plant science adapted by e.g. using suggestions from the Minimum Information About a Plant Phenotyping Experiment (MIAPPE).  
These unlike cross-domain minimal sets such as Dublin core, which mostly defines the submitter and what general type of data is being dealt with (e.g. images), allow reusability by other researchers as it also defines properties of the plant (see the preceding section). However, of course minimal cross-domain annotations are part of the  Example Project.    Other standards are also adhered to.  

Will search keywords be provided in the metadata to optimize the possibility for discovery and then potential re-use?

Keywords about the experiment and the general consortium will be included, as well as an abstract about the data, where useful. In addition, certain keywords can be auto-generated from dense metadata and its
underlying ontologies.

Will metadata be offered in such a way that it can be harvested and indexed?

To maintain data integrity and to be able to re-analyze data, data sets will get version numbers where this is useful (e.g. raw data must not be changed and will not get a version number and is considered
immutable). 

Data variables will be allocated standard names. For example, genes, proteins and metabolites will be named according to approved nomenclature and conventions. These will also be linked to functional ontologies where possible. Datasets will also be named I a meaningful way to ensure readability by humans. Plant names will include traditional names, binomials, and all strain/cultivar/subspecies/variety identifiers.

#### 2.2.    Making data accessible

Repository

Will the data be deposited in a trusted repository?

Data will be made available via the  Example Project  platform using a user-friendly front end that allows data visualization. Besides this it will be ensured that data which can be stored in
international discipline related repositories which use specialized technologies (Sequencing at the national US center Sequencing and sequence data at the EU center: EBI, Proteome database:
metabolomic database: ) will be used to store data and the data will be processed there as well.

Specialized repositories will be used where appropriate, such as INSDC (GenBank, EBI, DDBJ) for nucleotide sequence data, PIR/UniProt/SWISS-PROT for proteins, PDB for protein structures, GEO for transcriptomics, PRIDE for proteomics data, and METLIN for metabolomics data. For unstructured and less standardized data (e.g., experimental phenotypic measurements), these will be annotated with metadata and if complete allocated a digital object identifier (DOI).

Have you explored appropriate arrangements with the identified repository where your data will be deposited?

The submission is for free, and it is the goal (at least of ENA) to obtain as much data as possible. Therefore, arrangements are neither necessary nor useful. Catch-all repositories are not required.

Does the repository ensure that the data is assigned an identifier? Will the repository resolve the identifier to a digital object?

Data will be made available via the  Example Project  platform using a user-friendly front end that allows data visualization. Besides this it will be ensured that data which can be stored in
international discipline related repositories which use specialized technologies (Sequencing at the national US center Sequencing and sequence data at the EU center: EBI, Proteome database:
metabolomic database: ) will be used to store data and the data will be processed there as well. The ARC and the converters provided by DataPLANT will guarantee the upload into the endpoint repositories is fast and easy.

As noted above, specialized repositories like SRA /ENA, Pride /Proteomexchange are the most common ones and will be used when appropriate. In the case of unstructured less standardized data (e.g. experimental
phenotypic measurements), these will be metadata annotated and if complete given a digital object identifier (DOI).

Data:

Will all data be made openly available? If certain datasets cannot be shared (or need to be shared under restricted access conditions), explain why, clearly separating legal and contractual reasons from intentional restrictions. Note that in multi-beneficiary projects it is also possible for specific beneficiaries to keep their data closed if opening their data goes against their legitimate interests or other constraints as per the Grant Agreement.

By default, all data sets from the  Example Project  will be shared with the community and made openly available. However, before the data are released, all will be provided with an opportunity to check for potential IP (according to the consortium agreement and background IP rights). IP protection will be prioritized for datasets that offer the potential for exploitation.

Note that in multi-beneficiary projects it is also possible for specific beneficiaries to keep their data closed if relevant provisions are made in the consortium agreement and are in line with the reasons
for opting out.

If an embargo is applied to give time to publish or seek protection of the intellectual property (e.g. patents), specify why and how long this will apply, bearing in mind that research data should be made available as soon as possible.

The data will be published as soon as possible to guarantee reusability.   IP issues will be checked before publication.  All consortium partners will be
encouraged to make data available before publication, openly and/or under pre-publication agreements  such as those started in Fort Lauderdale and set forth by the Toronto International Data
Release Workshop.  This will be implemented as soon as IP-related checks are complete.

Will the data be accessible through a free and standardized access protocol?

Zenodo and the endpoint repositories will also be used for access. In General, web-based protocols are free and standardized for access.

If there are restrictions on use, how will access be provided to the data, both during and after the end of the project?

There are no restrictions, beyond the aforementioned IP checks, which are in line with e.g. European open data policies.

How will the identity of the person accessing the data be ascertained?

In case data is only shared within the consortium, if the data is not yet finished or under IP checks, the data is hosted internally and username and password will be required (see also our GDPR rules). In the case data is made public under final EU or US repositories, completely anonymous access is normally allowed. This is the case for ENA as well and both are in line with GDPR requirements.

Is there a need for a data access committee (e.g. to evaluate/approve access requests to personal/sensitive data)?

Consequently, there is no need for a committee.

Metadata:

Will metadata be made openly available and licenced under a public domain dedication CC0, as per the Grant Agreement? If not, please clarify why. Will metadata contain information to enable the user to access the data?

Yes, where possible, e.g. CC REL will be used for data not submitted to specialized repositories such as ENA.

How long will the data remain available and findable? Will metadata be guaranteed to remain available after data is no longer available?

The data will be made available for many years.
In any case data submitted to repositories (as detailed above) e.g. ENA /PRIDE would be subject to local data storage regulation.

Will documentation or reference about any software be needed to access or read the data be included? Will it be possible to include the relevant software (e.g. in open source code)?

No specialized software will be needed to access the data, usually just a modern browser. Access will be possible through web interfaces. For data processing after obtaining raw data, typical open-source software can be used. 
As stated above, here we use publicly available open-source and well-documented certified software

##### 2.3. Making data interoperable

What data and metadata vocabularies, standards, formats or methodologies will you follow to make your data interoperable to allow data exchange and re-use within and across disciplines? Will you follow community-endorsed interoperability best practices? Which ones?

As noted above, we foresee using minimal standards such as   MinSEQe for sequencing data and  Metabolights compatible forms for metabolites  and MIAPPE for phenotyping-like data . The minimal information standards will allow the integration of data across projects, and its reuse according to established and tested protocols.
Specialized repositories will be used for common data types. For unstructured and less standardized data (e.g., experimental phenotypic measurements), these will be annotated with metadata and if complete allocated a digital object identifier (DOI)..
Whenever possible, data will be stored in common and openly defined formats including all the necessary metadata to interpret and analyze data in a biological context. By default, no proprietary formats will be used. However Microsoft Excel files (according to ISO/IEC 29500-1:2016) might be used as intermediates by the consortium. In addition, text files might be edited in text processor files, but will be shared as pdf.
Open ontologies will be used where they are mature. As stated above, some ontologies and controlled vocabularies might need to be extended.

In case it is unavoidable that you use uncommon or generate project specific ontologies or vocabularies, will you provide mappings to more commonly used ontologies? Will you openly publish the generated ontologies or vocabularies to allow reusing, refining or extending them?

Common and open ontologies will be used. In fact, open biomedical ontologies will be used where they are mature. As stated in the previous question, sometimes ontologies and controlled vocabularies might have to be extended. . Ontology databases such as OBO Foundry will be used to publish ontology. .

Will your data include qualified references to other data (e.g. other data from your project, or datasets from previous research)?

The references to other data will be made in the form of DOI and ontology terms.

##### 2.4. Increase data re-use

How will you provide documentation needed to validate data analysis and facilitate data re-use (e.g. readme files with information on methodology, codebooks, data cleaning, analyses, variable definitions, units of measurement, etc.)?

The documentation will be provided in the form of ISA (Investigation Study Assay) and CWL (Common Workflow Language).

Will your data be made freely available in the public domain to permit the widest re-use possible? Will your data be licensed using standard reuse licenses, in line with the obligations set out in the Grant Agreement?

Yes, our data will be made freely available in the public domain to permit the widest re-use possible. Open licenses, such as Creative Commons (CC), will be used whenever possible.

Will the data produced in the project be useable by third parties, in particular after the end of the project?

There will be no restrictions once the data is made public.

Will the provenance of the data be thoroughly documented using the appropriate standards? Describe all relevant data quality assurance processes.

The  Example Project  has the following aim:  Example Aim.  Therefore, data collection, integration and visualization   through a standardized data management process is absolutely necessary  because the data are used not only to understand principles, but also be informed about the provenance of data analyzing data. Stakeholders must also be informed about the provenance of data. It is therefore necessary to ensure that the data are well generated and also well annotated with metadata using open standards, as laid out in the next section.

Describe all relevant data quality assurance processes. Further to the FAIR principles, DMPs should also address research outputs other than data, and should carefully consider aspects related to the allocation of resources, data security and ethical aspects.

The data will be checked and curated by using data collection protocol, personnel training, data cleaning, data analysis, and quality control . Document all data quality assurance processes, including the data collection protocol, data cleaning procedures, data analysis techniques, and quality control measures. This documentation should be kept for future reference and should be made available to stakeholders upon request.

#### 3    Other research outputs

In addition to the management of data, beneficiaries should also consider and plan for the management of other research outputs that may be generated or re-used throughout their projects. Such outputs can be either digital (e.g. software, workflows, protocols, models, etc.) or physical (e.g. new materials, antibodies, reagents, samples, etc.).

In the current data management plan, any digital output including but not limited to software, workflows, protocols, models, documents, templates, notebooks are all treated as data. Therefore, all aforementioned digital objects are already described in detail. For the non-digital objects, the data management plan will be closely connected to the digitalisation of the physical objects.

Beneficiaries should consider which of the questions pertaining to FAIR data above, can apply to the management of other research outputs, and should strive to provide sufficient detail on how their research outputs will be managed and shared, or made available for re-use, in line with the FAIR principles.

Open licenses, such as Creative Commons CC, will be used whenever possible even on the other digital objects.

#### 4.    Allocation of resources

What will the costs be for making data or other research outputs FAIR in your project (e.g. direct and indirect costs related to storage, archiving, re-use, security, etc.)?

The  Example Project  will bear the costs of data curation, and data maintenance/security before transfer to public repositories. Subsequent costs are then borne by the operators of these repositories.

Additionally, costs for after publication storage are incurred by end-point repositories (e.g. ENA) but not charged against the  Example Project  or its members but by the operation budget of these repositories.

How will these be covered? Note that costs related to research data/output management are eligible as part of the Horizon Europe grant (if compliant with the Grant Agreement conditions)

The cost born by the  Example Project  are covered by the project funding. Pre-existing structures will also be used.

Who will be responsible for data management in your project?

The responsible person will be  Example data officer name  of the  Example Project. 

How will long term preservation be ensured? Discuss the necessary resources to accomplish this (costs and potential value, who decides and how, what data will be kept and for how long)?

The data officer will ultimately decides on the strategy to preserve data that are not submitted to end-point subject area repositories when the
project ends. This will be in line with EU guidlines, institute policies, and data sharing based on EU and international standards.

#### 5.    Data security

What provisions are or will be in place for data security (including data recovery as well as secure storage/archiving and transfer of sensitive data)?

Online platforms will be protected by vulnerability scanning, two-factor authorization and daily automatic backups allowing immediate recovery. All partners holding confidential project data to use secure platforms with automatic backups and offsite secure copies.

Will the data be safely stored in trusted repositories for long term preservation and curation?

Wherever there are certified repositories, these will be used as end-point repositories.  Transcriptomics data and gene sequence data will be also made available upon publication via the standards
ENA/SRA,   metabolite data in e.g. Metabolights (and/or Nationwide repositories like the German NFDI the French INRAe),   Proteomics data in
e.g. Pride/Proteomexchange . In addition, the national resource will maintain safekeeping of data also after the project ends.

#### 6.    Ethics

Are there, or could there be, any ethics or legal issues that can have an impact on data sharing? These can also be discussed in the context of the ethics review. If relevant, include references to ethics deliverables and ethics chapter in the Description of the Action (DoA).

At the moment, we do not anticipate ethical or legal issues with data sharing. In terms of ethics, since this is plant data, there is no need for an ethics committee, however, diligence for plant resource benefit
sharing is considered (🡺see Nagoya protocol). <--pay attention to the sentence before because You have to check here and enter any due diligence here at the moment we are awaiting if Nagoya gets also part of sequence information. In any case
if you use material not from your (partner) country and characterize this physically e.g., metabolites, proteome, biochemically RNASeq etc. this might represent a Nagoya relevant action unless this is from e.g.
US (non partner), Ireland (not signed still contact them) etc but other laws might apply….

Will informed consent for data sharing and long term preservation be included in questionnaires dealing with personal data?

The only personal data that will potentially be stored is the submitter name and affiliation in the metadata for data. In addition, personal data will be collected for dissemination and communication
activities using specific methods and procedures developed by the  Example Project  partners to adhere to data protection. <--pay attention to the sentence before because You need to inform and better get WRITTEN consent that you store emails and
names or even pseudonyms such as twitter handles, we are very sorry about these issues we didn’t invent them

#### 7.    Other issues

Do you, or will you, make use of other national/funder/sectorial/departmental procedures for data management? If yes, which ones (please list and briefly describe them)?

Yes, the  Example Project  will use common Research Data Management (RDM) tools.

### 3     Annexes

#### 3.1     Abbreviations

CC Creative Commons

CC CEL Creative Commons Rights Expression Language

DDBJ DNA Data Bank of Japan

DMP Data Management Plan

DoA Description of Action

DOI Digital Object Identifier

EBI European Bioinformatics Institute

ENA European Nucleotide Archive

EU European Union

FAIR Findable Accessible Interoperable Reproducible

GDPR General data protection regulation (of the EU)

IP Intellectual Property

ISO International Organization for Standardization

MIAMET Minimal Information about Metabolite experiment

MIAPPE Minimal Information about Plant Phenotyping Experiment

MinSEQe Minimum Information about a high-throughput Sequencing Experiment

NCBI National Center for Biotechnology Information

NFDI National Research Data Infrastructure (of Germany)

NGS Next Generation Sequencing

RDM Research Data Management

RNASeq RNA Sequencing

SOP Standard Operating Procedures

SRA Short Read Archive

ONP Oxford Nanopore

qRTPCR quantitative real time polymerase chain reaction

WP Work Package
