## Supplementary material for "DataPLAN: a web-based data management plan generator for the plant sciences": Document_S3.doc

Export HTML To Doc

Data Management Plan of the DFG Project  Example Project

#### 1.    Data description

##### 1.1    Introduction

To best profit from open data, it is necessary not only to store data but to make data Findable, Accessible, Interoperable and Reusable (FAIR).

The aim of this document is to provide guidelines on principles guiding the data management in the  Example Project  and what data will be stored by using the responses to the DFG Data Management Plan (DMP) checklist
to generate a DMP document.

The detailed DMP instructs how data will be handled during and after the project. The  Example Project  DMP is modified according to the DFG data management checklist.

1.2    How does your project generate new data?

Data of different types or of different domains will be generated differently. For example:

- Short read sequencing will be either collected or outsourced and raw data will be received.
 - Metabolomic data will be generated using chromatography coupled to mass spectrometry and from enzyme platforms mostly.
 - Proteomic data will be generated using an EU platform which are in line with community standards.
- Image data will be generated by using equipment (cameras, scanners, and microscopes) or software. Original images which contain metadata such as exif photo information will be archived.
- Genomic data will be created from sequencing data. The sequencing data will be collected by Next Generation Sequencing (NGS) equipment. Then the sequencing data will be processed to get the genomic data.
- Genetic data will be generated by using Next Generation Sequencing (NGS) equipment.
- Targeted assays (e.g. glucose and fructose content) will be generated using specific equipment or experiments. The procedure is fully documented in the lab book.
- Models data will be generated by software simulations. The complete workflow, which includes the environment, runtime, parameter and results will be documented and achieved.
- The code data will be generated by programmers.
- The Excel data will be generated by experimentalists or data analysts by using Office or open-source software.
- The cloned DNA data will be generated by using a sequencing tool.
- Phenotypic data will be generated using phenotyping platforms.

The  Example Project  has the following aim:  Example Aim.  Therefore, data collection, integration and visualization   through a standardized data management process is absolutely necessary  because the data are used not only to understand principles, but also be informed about the provenance of data analyzing data. Stakeholders must also be informed about the provenance of data. It is therefore necessary to ensure that the data are well generated and also well annotated with metadata using open standards, as laid out in the next section.

Public data will be extracted as described in paragraph 1.3. For the  Example Project, specific data sets will be generated by the consortium partners.

1.3    Is existing data reused?

The project builds on existing data sets and relies on them.  For instance, without a proper genomic reference it is very difficult to analyze NGS data sets. It is also important to include existing data sets on the expression and metabolic behaviour of  Example Topic, but of course, also on existing characterization and the background knowledge. Genomic references can simply be gathered from reference databases for genomes/sequences, like the National Center for Biotechnology Information: NCBI (US); European Bioinformatics Institute: EBI (EU); DNA Data Bank of Japan: DDBJ (JP). Furthermore, prior 'unstructured' data in the form of publications and data contained therein will be used for decision making.

1.4    Which data types (in terms of data formats like image data, text data or measurement data) arise in your project and in what way are they further processed?

We foresee that the following data about  Example Topic  will be collected and generated at the very least: phenotypes, genotypes, other NGS data, metabolome, RNA-Seq and other forms of transcriptomic data, image datasets, proteome, targeted assays (e.g. glucose and fructose content),
model outputs, computational code, excel data, cloned DNA and result data. Furthermore, data derived from the original raw data sets will also be collected. This is important, as different analytical pipelines
might yield different results or include
ad-hoc

data analysis parts. Therefore, specific care will be taken, to document and archive these resources (including the
analytic pipelines) as well.

1.5    To what extent do these arise or what is the anticipated data volume?

We expect to generate raw data in the range of  ???  GB of data. The size of the derived data will be about  ???  GB.

#### 2.    Documentation and data quality

##### 2.1.    What approaches are being taken to describe the data in a comprehensible manner (such as the use of available metadata, documentation standards or ontologies)?

As noted above, we foresee using minimal standards such as   MinSEQe for sequencing data and  Metabolights compatible forms for metabolites  and MIAPPE for phenotyping-like data . The minimal information standards will allow the integration of data across projects, and its reuse according to established and tested protocols.
Specialized repositories will be used for common data types. For unstructured and less standardized data (e.g., experimental phenotypic measurements), these will be annotated with metadata and if complete allocated a digital object identifier (DOI)..
Whenever possible, data will be stored in common and openly defined formats including all the necessary metadata to interpret and analyze data in a biological context. By default, no proprietary formats will be used. However Microsoft Excel files (according to ISO/IEC 29500-1:2016) might be used as intermediates by the consortium. In addition, text files might be edited in text processor files, but will be shared as pdf.

We will use Investigation, Study, Assay (ISA) specification for metadata creation. For specific data (e.g., RNASeq or genomic data), we use metadata templates from the end-point repositories.  The Minimum Information About a Next-generation Sequencing Experiment (MinSEQe) will also be used. 

  Metabolights submission compliant standards will be used for metabolomic data where this is acccepted by the consortium partners.<--pay attention to the sentence before because some Metabolomics partners considers Metabolights
not an accepted standard As a part of plant research community, we use  MIAPPE for phenotyping data in the broadest sense, but we will also be rely on  specific SOPs for additional
annotations

#en

Keywords about the experiment and the general consortium will be included, as well as an abstract about the data, where useful. In addition, certain keywords can be auto-generated from dense metadata and
its underlying ontologies.

In fact, open biomedical ontologies will be used where they are mature. As stated in the previous question, sometimes ontologies and controlled vocabularies might have to be extended.

##### 2.2    What measures are being adopted to ensure high data quality?

The  Example Project  aims at the following aim:  Example Aim.  Therefore, data collection, integration and visualization   through a standardized data management process is absolutely necessary  because the data are used not only to understand principles, but also be informed about the provenance of data analyzing data. Stakeholders must also be informed about the provenance of data. It is therefore necessary to ensure that the data are well generated and also well annotated with metadata using open standards.
Data variables will be allocated standard names. For example, genes, proteins and metabolites will be named according to approved nomenclature and conventions. These will also be linked to functional ontologies where possible. Datasets will also be named I a meaningful way to ensure readability by humans. Plant names will include traditional names, binomials, and all strain/cultivar/subspecies/variety identifiers.

To maintain data integrity and to be able to re-analyze data, data sets will get version numbers where this is useful (e.g. raw data must not be changed and will not get a version number and is considered
immutable).

As mentioned above, we foresee using e.g.  MinSEQe for sequencing data and  Metabolights compatible forms for
metabolites as well as MIAPPE for phenotyping-like data. The latter will thus allow the integration of data across projects and safeguards that reuse established and tested protocols.
Additionally, we will use ontology terms to enrich the data sets relying on free and open ontologies. In addition, additional ontology terms might be created and be canonized during the  Example Project. 

##### 2.3    Are quality controls in place and if so, how do they operate?

The data will be checked and curated through the project period. Phd students and lab professionals will be responsible for the first-hand quality control. Afterwards, the data will be checked and annotated by  Example data officer name.   FastQC will be conducted on the base-calling. #enBefore publication, the data will be controlled again.

##### 2.4    Which digital methods and tools (e.g. software) are required to use the data?

The  Example Project  will use common Research Data Management (RDM) tools.

No specialized software will be needed to access the data, usually just a modern browser. Access will be possible through web interfaces. For data processing after obtaining raw data,
typical open-source software can be used. As no proprietary software is needed, no documentation needs to be provided.

As stated above, here we use publicly available open-source and well-documented certified software .

#### 3.    Storage and technical archiving the project

##### 3.1    How is the data to be stored and archived throughout the project duration?

Wherever there are certified repositories, these will be used as end-point repositories.  Transcriptomics data and gene sequence data will be also made available upon publication via the standards
ENA/SRA,   metabolite data in e.g. Metabolights (and/or Nationwide repositories like the German NFDI the French INRAe),   Proteomics data in
e.g. Pride/Proteomexchange . In addition, the national resource will maintain safekeeping of data also after the project ends.

Data will be made available for many years.

In any case data submitted to international
discipline related repositories which use specialized technologies (as detailed above) e.g. ENA /Pride would be subject to local data storage regulation.

##### 3.2    What is in place to secure sensitive data throughout the project duration (access and usage rights)?

In case data is only shared within the consortium, if the data is not yet finished or under IP checks, the data is hosted internally, and the username and the password will be required (see also our GDPR rules).
In the case data is made public under final EU or US repositories, completely anonymous access is normally allowed. this is the case for ENA as well and both are in line with GDPR requirements.

There will be no restrictions once the data is made public.

#### 4.    Legal obligations and conditions

##### 4.1    What are the legal specifics associated with the handling of research data in your project?

At the moment, we do not anticipate ethical or legal issues with data sharing. In terms of ethics, since this is plant data, there is no need for an ethics committee, however, diligence for plant resource
benefit sharing is considered (🡺see Nagoya protocol). <--pay attention to the sentence before because You have to check here and enter any due diligence here at the moment we are awaiting if Nagoya gets also part of sequence information.
In any case if you use material not from your (partner) country and characterize this physically e.g., metabolites, proteome, biochemically RNASeq etc. this might represent a Nagoya relevant action unless
this is from e.g. US (non partner), Ireland (not signed still contact them) etc but other laws might apply….

The only personal data that will potentially be stored is the submitter name and affiliation in the metadata for data. In addition, personal data will be collected for dissemination and communication
activities using specific methods and procedures developed by the  Example Project  partners to adhere to data protection. <--pay attention to the sentence before because You need to inform and better get WRITTEN consent that you store emails and
names or even pseudonyms such as twitter handles, we are very sorry about these issues we didn’t invent them

##### 4.2    Do you anticipate any implications or restrictions regarding subsequent publication or accessibility?

Once data is transferred to the  Example Project  platform, data security will be imposed. This comprises secure storage, and the use of
passwords and usernames is generally transferred via separate safe media.

##### 4.3    What is in place to consider aspects of use and copyright law as well as ownership issues?

Open licenses, such as Creative Commons (CC), will be used whenever possible.

##### 4.4    Are there any significant research codes or professional standards to be taken into account?

Whenever possible, data will be stored in common and openly defined formats including all the necessary metadata to interpret and analyze data in a biological context. By default, no proprietary formats will
be used; however, Microsoft Excel files (according to ISO/IEC 29500-1:2016) might be used as intermediates by the consortium. In addition,
text files might be edited in text processor files, but will be shared as pdf.

#### 5.    Data exchange and long-term data accessibility

##### 5.1    Which data sets are especially suitable for use in other contexts?

The data will be useful for the  Example Project  partners, the scientific community working on  Example Topic  or the general public interested in  Example Topic.  Hence, the  Example Project  also strives to collect the data
that has been disseminated and potentially advertise it, if it is not included in a publication anyway, which is the most
likely form of dissemination.

##### 5.2    Which criteria are used to select research data to make it available for subsequent use by others?

By default, all data sets from the  Example Project  will be shared with the community and made openly available. This is, however, after partners have had the ability to check for IP protection (according to
agreements and background rights). However, all partners also strive for IP protection of data sets which will
be tested and due diligence will be given.

Note that in multi-beneficiary projects it is also possible for specific beneficiaries to keep their data closed if relevant provisions are made in the consortium agreement and are in line with the
reasons for opting out.

##### 5.3    Are you planning to archive your data in a suitable infrastructure?

Data will be made available via the  Example Project  platform using a user-friendly front end that allows data visualization. Besides this it will be ensured that data which can be stored in
international discipline related repositories which use specialized technologies (Sequencing at the national US center Sequencing and sequence data at the EU center: EBI, Proteome database:
metabolomic database: ) will be used to store data and the data will be processed there as well. The ARC and the converters provided by DataPLANT will guarantee the upload into the endpoint repositories is fast and easy.

As noted above, specialized repositories like SRA /ENA, Pride /Proteomexchange are the most common ones and will be used when appropriate. In the case of unstructured less standardized data (e.g. experimental
phenotypic measurements), these will be metadata annotated and if complete given a digital object identifier (DOI).

The submission is for free, and it is the goal (at least of ENA) to obtain as much data as possible. Therefore, arrangements are neither necessary nor useful. Catch-all repositories are not required.

##### 5.4    If so, how and where? Are there any retention periods?

There are no restrictions, beyond the aforementioned IP checks, which are in line with e.g. European open data policies.

The decides on preservation of data not submitted to end-point subject area repositories after project end. This will be in line with EU
institute policies and data sharing based on EU and international standards.

##### 5.5    When is the research data available for use by third parties?

The data will be published as soon as possible to guarantee reusability.   In general, IP issues will first be checked.  All consortium partners will be
encouraged to make data available prior to publication openly and/or under pre-publication agreements  such as those started in Fort Lauderdale and set forth by the Toronto International Data
Release Workshop.

#### 6.    Responsibilities and resources

##### 6.1    Who is responsible for adequate handling of the research data (description of roles and responsibilities within the project)?

The responsible will be  Example data officer name  as data Officer.
The data responsible(s) (data officer) decides on the preservation of data not submitted to end-point subject area repositories after the
project end. This will be in line with EU institute policies, and data sharing based on EU and international standards.

##### 6.2    Which resources (costs; time or other) are required to implement adequate handling of research data within the project?

The costs comprise data curation, and maintenance on the  Example Project´ s side.

Additionally, last-level costs for storage are incurred by end-point repositories (e.g. ENA) but not charged against the  Example Project  or its members but by the operation budget of these repositories.

A large part of the cost is covered by the  Example Project  

##### 6.3    Who is responsible for curating the data once the project has ended?

As applicable,  Example data officer name, who is responsible for ongoing data maintenance will also take care of it after the finish of the  Example Project. 

### 7     Annexes

#### 7.1     Abbreviations

CC Creative Commons

CC CEL Creative Commons Rights Expression Language

DDBJ DNA Data Bank of Japan

DMP Data Management Plan

DoA Description of Action

DOI Digital Object Identifier

EBI European Bioinformatics Institute

ENA European Nucleotide Archive

EU European Union

FAIR Findable Accessible Interoperable Reproducible

GDPR General data protection regulation (of the EU)

IP Intellectual Property

ISO International Organization for Standardization

MIAMET Minimal Information about Metabolite experiment

MIAPPE Minimal Information about Plant Phenotyping Experiment

MinSEQe Minimum Information about a high-throughput Sequencing Experiment

NCBI National Center for Biotechnology Information

NFDI National Research Data Infrastructure (of Germany)

NGS Next Generation Sequencing

RDM Research Data Management

RNASeq RNA Sequencing

SOP Standard Operating Procedures

SRA Short Read Archive

ONP Oxford Nanopore

qRTPCR quantitative real time polymerase chain reaction

WP Work Package
