## Supplementary material for "DataPLAN: a web-based data management plan generator for the plant sciences": Document_S4.doc

Export HTML To Doc

### Practical Data Management Guide of the  Example Project

### **This practical guide of data management in the  Example Project  should be considered as a minimum description, leaving flexibility to include additional actions of specific domain or to national or local legislation.  The practical guide of data management in the  Example Project  aims at providing a complete walkthrough for the researcher. The contents are customized based on the user input in the Data Management Plant Generator (DMPG). The practices in this guide are customized to fit related legal, ethical, standardization and funding body requirements. The suitable practices will cover all steps of a data management life-cycle: 1. Data acquisition: 1. Data generation Data should be generated by devices that are compatible with the open-source format. The  Example Topic  should be compliant to biodiversity protocols. The protocols used to collect phenotypes, genotypes, other NGS data, metabolome, RNA-Seq and other forms of transcriptomic data data about  Example Topic  will be stored in a FAIR data storage.   2. Data collection The data collection process is conducted by experimental scientists and stewarded by  Example data officer name.   3. Data Organization The data organization process is conducted by  Example data officer name.  The detailed organization method and procedure are reported to the PIs.   2. Annotation 1. Workflow documentation Because the data collection process is conducted by experimental scientists and stewarded by  Example data officer name.   2. Metadata completion In case some of the metadata is still missing from the documentation from the experimental scientists and data officer. We foresee using e.g. MinSEQe for sequencing data and Metabolights compatible forms for metabolites as well as MIAPPE for phenotyping like data. The latter will thus allow the integration of data across projects and safeguards that reuse established and tested protocols. Additionally, we will use ontology terms to enrich the data sets relying on free and open ontologies. In addition, additional ontology terms might be created and be canonized during the  Example Project.  3. Maintenance: 1. Data storage Raw data collected in previous steps are stored immediately by using in a secure infrastructure. ARC (Annotated Research Context) is used as a container to store the raw data as well as metadata and workflow. 2. Data curation Data is curated regularly as long as there are needs for update or revision. 4. Publication and sharing 1. Data publishing Transcriptomics data and gene sequence data will be also made available upon publication via the standards ENA/SRA. Metabolite data in e.g. Metabolights (and/or Nationwide repositories like the German NFDI the French INRAe). and Proteomics data in e.g. Pride/Proteomexchange. In addition, the national resource will maintain safekeeping of data also after the project ends. 2. Data sharing In case data is only shared within the consortium, if the data is not yet finished or under IP checks, the data is hosted internally, and the username and the password will be required (see also our GDPR rules). In the case data is made public under final EU or US repositories, completely anonymous access is normally allowed. This is the case for ENA as well and both are in line with GDPR requirements.**

##### Metadata focus timeline

### **| Stages | Actions | | --- | --- | | Study initialization | The metadata of study is created at the beginning of the project and updated continuously afterwards | | Sample Collection | The information used to identify exact samples are initiated before experiments and updated at assay creation stages. | | Assay Creation | Assay metadata must be collected as a daily routine during the experimental phrase. | | Computational Analysis | Workflow annotation will be conducted during the computational analysis phrase. | | Results Sharing | The metadata of results are collected after all modifications and should not be changed after publication. |**

##### Preferred formats for raw data

### | | | | --- | --- | | extension\_ident | Format Name | | .h5 | Hierarchical Data Format | | .bam | compressed binary version of a SAM file | | .cram | compressed columnar file format for storing biological sequences aligned to a reference sequence | | .fa | fasta | | .faa | fasta | | .fas | fasta | | .fasta | fasta | | .fastq | fastq | | .ffn | fasta | | .fna | fasta | | .fq | fastq | | .frn | fasta | | .sff | sff-trim | | | | | --- | --- | | .bam | compressed binary version of a SAM file | | .cram | compressed columnar file format for storing biological sequences aligned to a reference sequence | | .fa | fasta | | .faa | fasta | | .fas | fasta | | .fast5 | HDF5 | | .fasta | fasta | | .fastq | fastq | | .ffn | fasta | | .fna | fasta | | .fq | fastq | | .frn | fasta | | .sff | sff-trim | | bas.h5 | HDF5 | | .h5 | Hierarchical Data Format |   | | | | --- | --- | | .cdf | netCDF (AIA/ANDI) interchange data format | | .cmp | netCDF compare file | | .abf | Axon Binary File | | .d | Agilent | | .dat | Chromtech, Finnigan, VG | | .idb | MASSLAB binary file | | .jpf | Mass Center Main Mass Spectrometry Data (JEOL USA, Inc.) | | .lcd | Shimadzu LC Solution / Labsolutions Data File | | .mgf | Mascot Generic File | | .raw | Thermo Xcalibur, Micromass (Waters), PerkinElmer, Waters | | .scan | a spectrum or a Total Ion Chromatogram (TIC) | | .wiff | ABI/Sciex | | .xps | Thermo Fisher Scientific K-Alpha+ spectrometer file | | cdf.cmp | netCDF compare file |   | | | | --- | --- | | .baf | Bruker | | .d | Agilent | | .dat | Chromtech, Finnigan, VG | | .fid | Bruker | | .ita | ION-TOF | | .itm | ION-TOF | | .mgf | Mascot Generic File | | .ms | Finnigan (Thermo) | | .ms2 | Sequest MS/MS peak list | | .pkl | Micromass peak list | | .qgd | Shimadzu | | .qgd | Shimadzu | | .raw | Thermo Xcalibur, Micromass (Waters), PerkinElmer, Waters | | .raw | Physical Electronics/ULVAC-PHI | | .sms | Bruker/Varian | | .spc | Shimadzu | | .splib | spectral library file | | .t2d | ABI/Sciex | | .tdc | Physical Electronics/ULVAC-PHI | | .wiff | ABI/Sciex | | .xms | Bruker/Varian | | .yep | Bruker | | .dta | Sequest MS/MS peak list | | .msp | | | .nist | |
